# Targeted next-generation sequencing of Candidate Regions Identified by GWAS Revealed SNPs Associated with IBD in GSDs

**DOI:** 10.1101/2021.04.20.440584

**Authors:** Atiyeh Peiravan, Mazdak Salavati, Androniki Psifidi, Mellora Sharman, Andrew Kent, Penny Watson, Karin Allenspach, Dirk Werling

## Abstract

Canine Inflammatory bowel disease (IBD) is a chronic multifactorial disease, resulting from complex interactions between the intestinal immune system, microbiota and environmental factors in genetically predisposed dogs. Previously, we identified several single nucleotide polymorphisms (SNP) and regions on chromosomes (Chr) 7, 9, 11 and 13 associated with IBD in German shepherd dogs (GSD) using GWAS and FST association analyses. Here, building on our previous results, we performed a targeted next-generation sequencing (NGS) of a two Mb region on Chr 9 and 11 that included 14 of the newly identified candidate genes, in order to identify potential functional SNPs that could explain these association signals. Furthermore, correlations between genotype and treatment response were estimated. Results revealed several SNPs in the genes for canine *EEF1A1*, *MDH2*, *IL3*, *IL4*, *IL13* and *PDLIM*, which, based on the known function of their corresponding proteins, further our insight into the pathogenesis of IBD in dogs. In addition, several pathways involved in innate and adaptive immunity and inflammatory responses (i.e. T helper cell differentiation, Th1 and Th2 activation pathway, communication between innate and adaptive immune cells and differential regulation of cytokine production in intestinal epithelial cells by IL-17A and IL-17F), were constructed involving the gene products in the candidate regions for IBD susceptibility. Interestingly, some of the identified SNPs were present in only one outcome group, suggesting that different genetic factors are involved in the pathogenesis of IBD in different treatment response groups. This also highlights potential genetic markers to predict the response in dogs treated for IBD.

## Introduction

Inflammatory bowel disease (IBD) is a common cause of chronic gastrointestinal disease in humans and dogs. IBD is characterised by persistent or recurrent gastrointestinal signs (GI) including chronic diarrhoea, vomiting and weight loss and with histological evidence of inflammation in the lamina propria of the small intestine, large intestine or both (Albert 1999). The diagnosis of IBD in both humans and dogs is by exclusion, as several diseases can cause chronic gastrointestinal inflammation secondarily (Guilford 1996; Hendrickson et al., 2002).

The pathogenesis of IBD is believed to be multifactorial in both humans and dogs, caused by a complex interaction between the intestinal innate and adaptive immune system, the intestinal microbiome, and the genetic make-up of an individual. Although IBD can affect multiple dog breeds, breed-specific disease phenotypes and associations have been reported (Kimmel et al., 2000; Simpson et al. 2006). In the United Kingdom (UK), German shepherd dogs (GSD) have been reported to be at increased risk of developing the disease (Kathrani et al., 2011).

IBD is not a curable disease, therefore the aim of current treatment approches is to minimise the severity and frequency of the clinical signs. In general, treatment protocols include dietary modifications (Luckschander et al. 2006; Mandigers et al. 2010), antibiotics (German et al., 2003; Westermarck et al., 2005), and corticosteroid (Allenspach et al., 2007) treatment trials. As such, canine IBD is generally classified only retrospectively based on response to treatment into food responsive disease/diarrhoea (FRD), antibiotic responsive disease/diarrhoea (ARD), or steroid responsive disease/diarrhoea (SRD), which are usually used interchangeably with idiopathic IBD. Canine IBD patients are considered as FRD and ARD if their clinical signs improve or resolve following dietary modification and/ or antibiotic treatment. Those that fail to respond to a change of diet and/ or antibiotic therapy require immunosuppressive treatment (usually corticosteroids) to treat their clinical signs (SRD) (Allenspach et al., 2007). To date, treatment with anti-inflammatory/ immunomodulatory medication such as corticosteroids is the mainstay treatment for both human and canine IBD patients (German et al., 2003). However, up to 50% of dogs with IBD that are initially managed with steroids will develop resistance and/or significant side effects, which ultimately leads to euthanasia for many of them (Allenspach et al., 2006, 2007).

Recent advances in clinical genetics make it possible to use the patients’ genetic profile to predict response to treatment (William and Sandborn 2004; Roberts and Barclay 2012). Similar to human IBD, it is hoped that identifiyng the genes involved in canine IBD will provide insights into disease pathogenesis in canine IBD. This could lead to the development of genetic screening panels useful for both diagnosis and identifying dogs that are more likely to fall into specific groups of treatments.

So far, studies of canine IBD using a candidate gene approach, have identified a number of single nucleotide polymorphisms (SNP) associated with the disease in genes encoding pattern recognision receptors (PRR) of the innate immunity (Kathrani et al., 2010, 2011, 2014) as well as associations between SNPs in Major histocompatibility class (MHC) II haplotypes and a potentially increased resistance to IBD in GSD (Peiravan et al., 2016). The release of the re-assembled dog genome and development of high-density canine DNA SNP arrays have enabled several successful GWAS studies aimed at investigating the genetic architecture of both monogenic and polygenic complex diseases of the dog (Wood et al., 2009; Wilbe et al., 2010; Lequarré et al., 2011; Tengvall et al., 2013). Previously, based on GWAS and FST association analyses of IBD cases and controls, we identified several SNPs and regions on chromosomes (Chr) 7,9,11 and 13 associated with IBD in GSD, including a total of 80 genes. Using a combination of pathways analysis and a list of genes that have been reported to be involved in human IBD, we identified 16 candidate genes potentially associated with IBD in GSD (Peiravan et al. 2018).

Genome-wide association studies rely on the principal of linkage disequilibrium (LD). While the extensive LD and long haplotype blocks (0.5-1.0 Mb) within breeds, resulted from genetic bottlenecks during domestication of dogs and breed formation, is an advantage in the initial GWAS it complicates the subsequent identification of the causative variant(s) (Sutter 2004; Karlsson et al., 2007). The association signals identified through GWAS represent most likely markers that are not the causal variants themselves but are linked instead with nearby causative polymorphisms. Therefore, in order to generate a hypothesis about mechanisms underlying a specific phenotype, it is important to detect the causal variants themselves.

In the present study, we performed a targeted NGS of 2 Mb regions on Chr 9 and 11, which include 14 of the newly identified candidate genes, aiming to identify potential functional SNPs that could explain the GWAS association signal. We also investigated whether there was a correlation between the identified SNPs and response to treatment in the IBD cases, used in this study.

## Materials and methods

### Ethics and welfare statement

All blood samples used in this study were collected in ethylenediaminetetraacetic acid (EDTA) and represented residual material following completion of clinical diagnostic testing. Residual samples were utilised for research with informed owner consent. The use of these residual EDTA blood samples and buccal swab samples within the current study was approved by the RVC Animal Welfare Ethical Review Body (AWERB; approval number 2013 1210).

### Selection of cases and controls for targeted sequencing

IBD cases and controls were identified based on inclusion criteria described previously (Peiravan et al., 2018). A follow-up on all cases and controls was performed by contacting the referring veterinary surgeons and/or owners to gather information on treatment response of the dogs, including assessment of any changes to the course of treatment, and if so what the response to the new treatment was.

A total number of 48 GSDs with adequate followup information and available genomic (g)DNA samples were enrolled, including 28 cases (diagnosed with IBD) and 20 controls (non-inflammatory disease). Among the IBD cases, there were 9 FRD, 4 ARD, 11 SRD and 4 NRS/PTS (No Response to Steroid, PTS: Put To Sleep) cases. Control dogs were breed-matched that were presented with variety of non-inflammatory conditions or no-known diseases.

### SureSelect XT Library preparation and sequence capture

A custom-designed sequence capture array (SelectSure XT custom 0.5-2.9 Mb, Agilent), was designed and manufactured by Agilent, in order to isolate the targeted region identified on Chr 9 and 11, 1 Mb up- and downstream of the most significant SNPs identified by the previous GWAS (Peiravan et al., 2018), from total gDNA, using start and end coordinates of the associated regions. Properties of the final design of each array designed for the capture of the target regions on Chr 9 and 11 are shown in Table 1.

**Figure 1.**
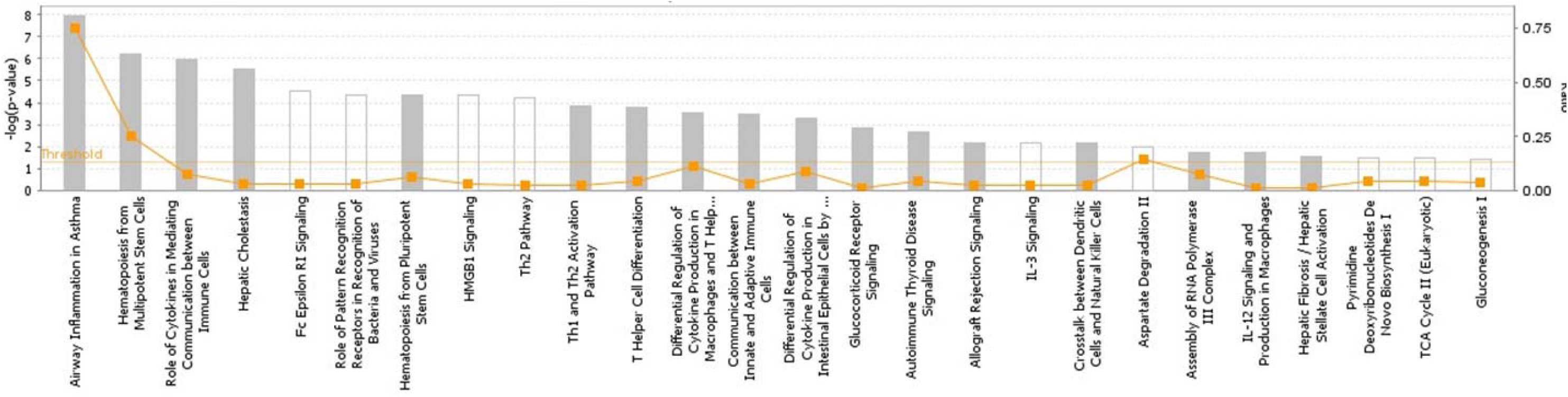
The most highly represented canonical pathways of genes located at the candidate regions. The solid yellow line indicates the significance threshold. The line with squares represents the ratio of the genes represented within each pathway to the total number of genes in the pathway.

**Figure 2.**
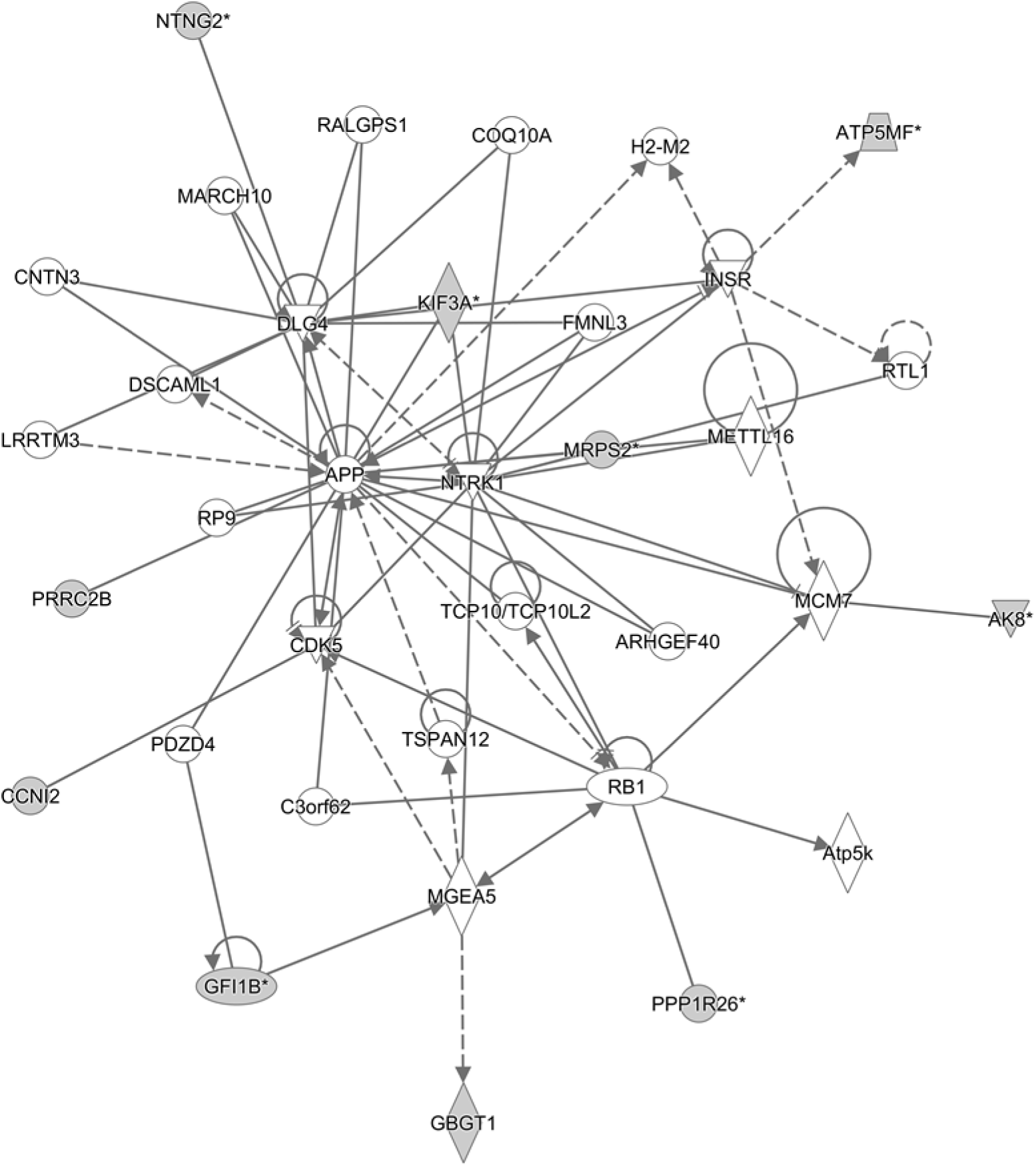
Cell cycle network and genes involved in the network. Grey filled shapes represent genes included in the list of candidate genes identified in the targeted regions. Solid and dotted lines represent direct and indirect interaction between genes respectively.

**Table 1.**
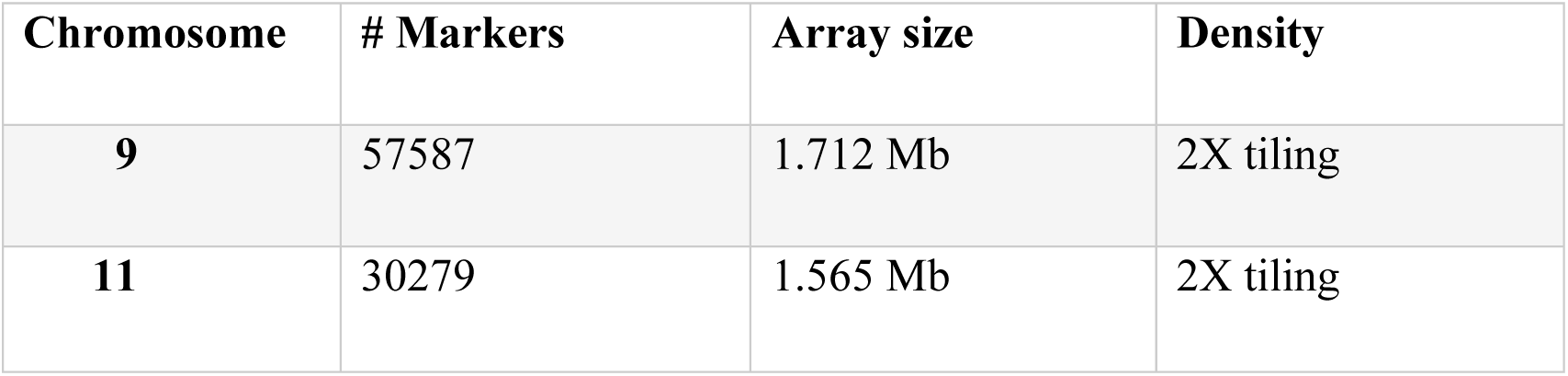
Properties of the capture arrayes designed for isolation of the target regions on Chromosomes 9 and 11.

For each gDNA sample to be sequenced, an individual indexed library was prepared. DNA libraries were prepared using the SureSelect XT Library Prep Kit (Agilent) following the manufacturer instructions. In brief, 200 ng of the gDNA samples were fragmented by sonication and the paired-end adaptors were ligated to the blunt-ended fragments using the SureSelect XT Library Prep Kit ILM. Then, the adaptor-ligated fragments were PCR amplified, purified and hybridyzed to the capture array, using the SureSelectXT DNA kit following, the manufacturer instructions. Then, unbound fragments were washed away. Subsequently, the SureSelect-captured DNA libraries were eluted, purified and PCR amplified using an individual indexing primer for each sample. Following quality control, captured libraries were sequenced with a read length of 150 bp (paired-end reads) on an Illumina NextSeq (500 Mid Output) platform. Sequencing was performed at the Genomic Centre of Queen Mary University of London (QMUL).

### Analysis of sequencing data

The Genome Analysis Toolkit (GATK 3.8) (https://software.broadinstitute.org/gatk) best practice workflow, in house bash and R scripting was used for processing of raw sequencing data. BAMStats 2.1, which provides descriptive statistics for various metrics, was used to calculate average coverage.

Raw sequencing data were visualised and inspected using FastQC (v0.11.6) (http://www.bioinformatics.babraham.ac.uk/projects/fastqc/). Thereafter, reads were trimmed and filtered using Trimmomatic (v0.36) (Bolger et al., 2014). The trimmed reads were aligned and mapped to the canine genome (CanFam3.1), using Burrow-wheeler aligner (BWA, v0.7.15) with default parameters (bwa mem -M -t -R). BWA– MEM is designed for aligning relatively short sequence reads ranging from 70 bp to 1 Mb against a long reference and is generally recommended for high-quality queries (Li and Durbin 2009). Once the reads were mapped to the canine genome and merged, they were sorted by the coordinates using the “samtools sort” command (Li and Durbin 2009).

Potencial PCR duplicates were flagged in the read’s SAM record using Picard tool (hosted by SAMtools), so that duplicates could be identified during downstream processing. Most GATK tools will then ignore the flagged reads by default, through the internal application of a read filter. Base quality score was adjusted according to a model of covariations bulit based on the data and a set of known variants (https://m.ensembl.org/info/genome/variation/species/sources_documentation.html#canis_lupus_familiaris).

Genetic variants were called individually on each sample’s BAM file(s) using the HaplotypeCaller (GATK 3.8) in -ERC gVCF mode to produce an intermediate file format called gVCF file (genomic VCF). Following variant calling, the GVCFs of multiple samples were run through joint genotyping to produce a multi-sample VCF callset, using GenotypeGVCFs. Then, the raw SNPs extracted from the multi-sample VCF callset produced at genotyping stage, was subject to hard filtering. All the scripts that were used for variant calling, genotyping, and filtering annotations and values recommended by GATK best practice for hard filtering are shown in Supplementary Table A1.

### Analysis of variants

The coordinate positions of the filtered SNPs were used to categorise them into different groups; a) SNPs only present in cases, b) SNPs only present in controls and c) SNPs present in both cases and controls. The latest group was further divided into SNPs with the same alternate allele in both populations and those with different alternate in case or control population.

SNPs in each group were then annotated using variant effect predictor (VEP, https://www.ensembl.org/Tools/VEP). The effect of SNPs on genes, transcripts, and protein sequence, as well as regulatory regions were determined and SIFT predictions of the effects of SNPs (tolerated or deleterious) were also acquired through VEP. The focus was on detection of exonic SNPs with high importance such as non-sense (stop-gain) and missense (non-synonymous) exonic and splicing, since these can affect considerably the function of the encoded protein. In addition, genomic regions located within 1 Kb up- and downstream of the candidate genes were analysed in order to detect SNPs with putative regulatory effect. For the case-control (overlapped) group with the same alternate allele in case and control group, high impact SNPs with deleterious SIFT score were detected and then the frequency of the identified SNPs copmpared between the two groups to identify those with statisticaly significant difference between cases and controls. The integrative genomic viewer (IGV 2.4.) was also used for manual visualisation of SNPs.

### Statistical analysis

Allele frequencies were calculated and compared using VassarStats (Web Site for Statistical Computation, Vassar College, Poughkeepsie, NY, http://www.vassarstats.net/odds2x2.html). Two-way contingency tables were used to calculate two-tailed Fisher’s exact probability statistic for association of each allele with disease status. Statistical significance was set at a p value of < 0.05. The calculation of Hardy-Weinberg equilibrium (HWE) for the identified variants was carried out performing chi-square test using the SNPSTATS programme (http://bioinfo.iconcologia.net/SNPstats).

### Pathway and enrichment analysis

The candidate gene lists for IBD susceptibility were analysed using the IPA programme (www.ingenuity.com) in order to identify canonical pathways and gene networks constructed by the products of the genes. IPA builds several possible pathways and networks serving as hypotheses for the biological mechanism underlying the data based on a large-scale causal network obtained from the Ingenuity Knowledge Base, which are subsequently summarised by the identification of the most suitable pathways and networks based on their statistical significance.

In addition to IPA analysis, a second approach was used to identify the best candidate genes associated with IBD in GSDs, using Enrichr, which is a web-based enrichment analysis tool (http://amp.pharm.mssm.edu/Enrichr). For this analysis, the default statistical tests and corrections for multiple testing to maintain an overall p value of 0.05 were applied.

### Association of variants with treatment response

To investigate whether there was a correlation between variants identified and response to treatment, the 28 IBD cases that were used in the current study were categorised into two groups as shown in Table 2.

**Table 2.** The number of cases in each treatment esponse group. FRD: food responsive disease/diarrhoea, ARD: antibiotic responsive disease/diarrhoea, SRD: steroid responsive disease/diarrhoea, NRS: no response to treatment, PTS: put to sleep (euthanized)

| Response to treatment |  | # cases |
| --- | --- | --- |
| Food and antibiotic treated cases | FRD | 9 |
|  | ARD | 4 |
| Steroid treated cases | SRD | 11 |
|  | NRS/PTS | 4 |

**Table 3.** Number of raw and hard filtered SNPs in the case and control groups in the target region on chromosomes 9 and 11.

| **Chromosome** | **Sample** | **Raw SNPs** | **Hard Filtered SNPs** |
| --- | --- | --- | --- |
| **9** | **Case** | 12149 | 10600 |
|  | **Control** | 12316 | 10908 |
| **11** | **Case** | 3806 | 2495 |
|  | **Control** | 3884 | 2508 |

**Table 4.** Number of SNPs in each group on Chr 9 and 11. Case only: SNPs that were only present in case population, Ctrl only: SNPs present only in control population, Overlapped with different Alt in Case/ or Ctrl: SNPs that were present in both case and control population but showing different alternate allele in case and control, Overlapped with same Alt in Case and Ctrl: SNPs that were present in both case and control population and also showing same alternate allele in both case and control. Chr: chromosome, Alt: alternate, Ctrl: control.

| **Chromosome** | **Case only** | **Control only** | **Overlapped with different Alt in Case** | **Overlapped with different Alt in Ctrl** | **Overlapped with same Alt in Case and Ctrl** |
| --- | --- | --- | --- | --- | --- |
| **9** | 985 | 1293 | 14 | 14 | 9601 |
| **11** | 721 | 734 | 6 | 6 | 1768 |

**Table 5.** Chr 9, variants with statistically significant differences in frequency between the case and control populations (present only in the case population). Hom Ref: sites with reference allele (AA), Het: Heterozygous (AB), Homo: Homozygous (BB), P value: Fisher exact probability test two tailed p value. COL5A1: Collagen Type V Alpha 1 Chain, MRPS2: Mitochondrial ribosomal protein subunit 2, EEF1A1: Eukaryotic elongation f1 lpha-1, MDH2: Malate dehydrogenase 2, NTNG2: Netrin-G2.

| Variation | Location | Allele | Consequence | Impact | Symbol | Case |  |  |  |  | Control |  |  |  |
| --- | --- | --- | --- | --- | --- | --- | --- | --- | --- | --- | --- | --- | --- | --- |
|  |  |  |  |  |  | No call | Hom Ref | Variant | Het | Homo | No call | Hom Ref | Variant | P value |
| . | 9:50769631 | G*/A | Upstream | MODIFIER | COL5A1 | 3 | 14 | 11 | 10 | 1 | 1 | 19 | 0 | 0.0009 |
| . | 9:51215957 | T*/C | downstream | MODIFIER | MRPS2 | 0 | 20 | 8 | 8 | 0 | 3 | 17 | 0 | 0.017 |
| **rs24555947** | 9:51227287 | C*/G | downstream | MODIFIER | EEF1A1 | 0 | 12 | 16 | 16 | 0 | 0 | 20 | 0 | 2E-05 |
| . | 9:51227325 | G*/A | downstream | MODIFIER | EEF1A1 | 3 | 6 | 19 | 19 | 0 | 2 | 18 | 0 | 6.68E-07 |
| . | 9:51227411 | T*/A | downstream | MODIFIER | EEF1A1 | 0 | 2 | 26 | 26 | 0 | 0 | 20 | 0 | 1.38E-11 |
| . | 9:51228186 | C*/T | stop_gained | HIGH | EEF1A1 | 0 | 2 | 26 | 26 | 0 | 0 | 20 | 0 | 1.38E-11 |
| **rs24570325** | 9:51228828 | G*/T | Upstream | MODIFIER | EEF1A1 | 9 | 12 | 7 | 4 | 3 | 0 | 20 | 0 | 0.003 |
| **rs24570326** | 9:51228829 | C*/T | Upstream | MODIFIER | EEF1A1 | 9 | 12 | 7 | 4 | 3 | 0 | 20 | 0 | 0.003 |
| . | 9:51255622 | T*/C | Upstream | MODIFIER | Novel gene | 0 | 17 | 11 | 9 | 2 | 0 | 20 | 0 | 0.001 |
| . | 9:51404724 | C*/T | Missense | MODERATE | MDH2 | 7 | 3 | 18 | 18 | 0 | 1 | 19 | 0 | 7.52E-09 |
| . | 9:51405189 | T*/C | downstream | MODIFIER | MDH2 | 4 | 1 | 23 | 23 | 0 | 1 | 19 | 0 | 2.50E-11 |
| **rs852815996** | 9:51994360 | T*/C | downstream | MODIFIER | NTNG2 | 8 | 8 | 12 | 2 | 10 | 3 | 17 | 0 | 0.00007 |

**Table 6.**
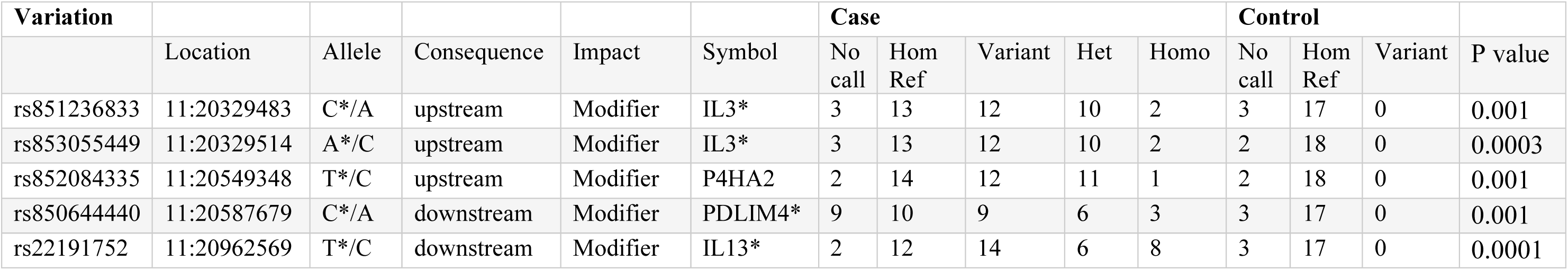
Chr 11, variants with statistically significant differences in frequency between the case and control populations (present only in the case population). Hom Ref: sites with reference allele (AA), Het: Heterozygous (AB), Homo: Homozygous (BB), P value: Fisher exact probability test two tailed p value. *IL3*: Interleukin 3, *P4HA2*: Prolyl 4-hydroxylase subunit alpha-2, *PDLIM4*: PDZ and LIM domain protein 4, *IL13*: Interleukine 13. *Genes already found to be associated with human IBD.

| Variation | Location | Allele | Consequence | Impact | Symbol | Case |  |  |  |  | Control |  |  | P value |
| --- | --- | --- | --- | --- | --- | --- | --- | --- | --- | --- | --- | --- | --- | --- |
|  |  |  |  |  |  | No call | Hom Ref | Variant | Het | Homo | No call | Hom Ref | Variant |  |
| rs851236833 | 11:20329483 | C*/A | upstream | Modifier | IL3* | 3 | 13 | 12 | 10 | 2 | 3 | 17 | 0 | 0.001 |
| rs853055449 | 11:20329514 | A*/C | upstream | Modifier | IL3* | 3 | 13 | 12 | 10 | 2 | 2 | 18 | 0 | 0.0003 |
| rs852084335 | 11:20549348 | T*/C | upstream | Modifier | P4HA2 | 2 | 14 | 12 | 11 | 1 | 2 | 18 | 0 | 0.001 |
| rs850644440 | 11:20587679 | C*/A | downstream | Modifier | PDLIM4* | 9 | 10 | 9 | 6 | 3 | 3 | 17 | 0 | 0.001 |
| rs22191752 | 11:20962569 | T*/C | downstream | Modifier | IL13* | 2 | 12 | 14 | 6 | 8 | 3 | 17 | 0 | 0.0001 |

**Table 7.**
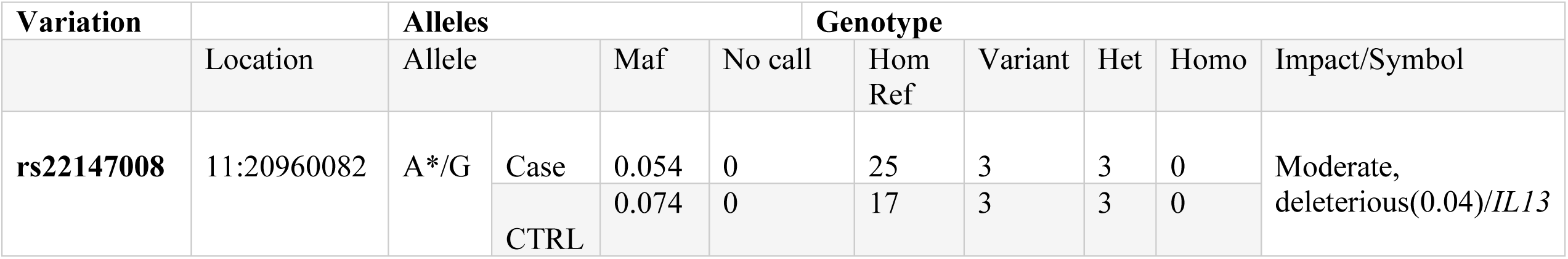
The only variant overlapped between the case and control populations, with same ALT in the case and control, with deleterious SIFT score.

## Results

### Quality control of gDNA, captured DNA libraries and sequencing data

The results of Qubit system, used to quantify gDNA before library preparation, are shown in Supplementary Table A2. The quality and size of fragmented DNA was assessed with D1000 ScreenTape on Agilent 2200 TapeStation. The electropherogram produced for each sample confirmed that the DNA samples had been fragmented to the required size with the shearing size range of 100 to 500 bp, with a peak at 150-200 bp. The concentration of the amplified DNA libraries after capture, the number of libraries and their size are shown in Supplementary Table A3. The electropherogram produced for each sample showed a distribution with a DNA fragment size peak of approximately 225 to 275 bp.

All samples were successfully sequenced and passed internal QC at the genomic centre of QMUL. The distribution of coverage was similar between the two target regions. Average mapped-sequencing coverage was 256× and 230× for 97.5% of the regions on Chr 9 and 11 respectively (Supplementary Table B). There was no significant differences in coverage rates between cases and controls on any of the two target regions. Average coverage of the target region on Chr 9 was 254× and 258× for cases and controls respectively. For the target region on Chr 11, the average coverage was 228× for cases and 233× for controls.

### Analysis of variants and variant annotation

The number of raw SNPs within the target regions on Chr 9 and 11 and the number of SNPs that passed hard filtering are shown in **Error! Reference source not found.** . **Error! Reference source not found.** shows the number of SNPs in each group per chromosome. SNPS identified in each group and their effect and properties are presented in the Supplementay Tables C and D.

### Variants on Chromosome 9

Twelve variants with high or moderate impact, which were deleterious based on their SIFT score, were detected only in cases (**Error! Reference source not found.Error! Reference source not found.**). Among these, there were two stop_ gained variants with high impact. The remaining ten variants were missense variants with moderate impact. In addition, 48 variants with modifier impact were identified within 1kb up- and downstream of the candidate genes present only in cases (Supplementary Table C).

Thirty missense variants (2 known and 28 new) with moderate impact that were deleterious according to their SIFT score were detected only in controls. In addition, 65 variants with modifier impact (13 known and 52 new) were observed within 1kb up- and downstream of genes (Supplementary Table C).

Twenty-six missense variants with moderate impact were identified (14 known and 12 new) in both cases and controls. However, no significant differences in the frequency of these variants was detected between cases and controls (Supplementary Table C).

### Variants on Chromosome 11

One new missense SNP (deleterious based on SIFT score, SIFT: 0.03) was found only in cases, and this SNP seems to have a moderate impact in a novel protein coding gene that is orthologous to human *ATP5MF* (ATP synthase membrane subunit F; for more details, please see Supplementary Table D). In addition, five SNPs (3 known and 2 new) were identified within 1kb upstream of gene TSS coordinates. In addition, 12 SNPs (3 known and 8 new) were detected within 1 Kb downstream of the hypothesised candidate genes. Interestingly, associations between several of these candidate genes and human IBD have already been described. The details of the identified variants are shown in Supplementary Table D.

Two of these SNPs with modifier impact were located within 1 kb upstream of *IL3*, a haemopoietic cytokine driving the development of myeloid stem cells that was previously identified to be associated with IBD (Peiravan et al. 2018). Two SNPs were found downstream of *PDLIM* (a protein with cysteine-rich double zinc fingers involved in protein-protein interaction and cytoskeletal organisation) and *IL13* (a Th2 cytokine involved in IgE synthesis, chitinase upregulation and hyperresponsiveness of mucosal surfaces) and one new SNP was found downstream of *IL4* (a Th2 cytokine produced by mast cells, eosinophils and basophils that stimulates B cells into plasma cells and shares functions similar to IL-13). All of these genes have already been reported to be associated with human IBD in previous studies. SNPs with a statistically significant difference (p value <0.05) between cases and controls are shown in **Error! Reference source not found.**6.

Two new missense SNPs, deleterious based on SIFT score, were identified in a protein coding gene, orthologous to human *HINT1* (histidine triad nucleotide binding protein 1) in the control population. HINT1 is a hydrolase and gene ontology annotations related to this gene include nucleotide binding and protein kinase C binding. Furthermore, five SNPs (1 known and 4 new) within 1 Kb upstream and ten SNPs (1 known and 9 new) within 1 Kb downstream of the candidate genes and Small nucleolar RNAs (snoRNA) on Chr 11 were identified in the control population. Two SNPs downstream of a snoRNA were observed to have significant P value (Supplementary Table D).

Ten missense SNPs with high and moderate impact were detected in both cases and controls. However, only one SNP in *IL13* was found to be deleterious based on the SFIT score and there was no statistically significant difference in the frequency of this SNP between case (Maf:0.054) and control group (Maf:0.074)(**Error! Reference source not found.**7).

### Pathway and Network analysis reveal impact on genes involved in inflammatory response and metabolism

#### IPA analysis

Several pathways involved in innate and adaptive immune and inflammatory response (i.e. T helper cell differentiation, Th1 and Th2 activation pathway, communication between innate and adaptive immune cells and differential regulation of cytokine production in intestinal epithelial cells by IL-17A and IL-17F) and metabolism (i.e. TCA cycle II (Eukaryotic), gluconeogenesis I, aspartate degradation II and pyrimidine deoxyribonucleotides De Novo biosynthesis) were constructed by the gene products in the candidate regions for IBD susceptibility (**Error! Reference source not found.** and Supplementary Table E). Moreover, four networks of molecular interactions related to cell cycle (IPA Score: 22) (**Error! Reference source not found.**), hereditary disorder and metabolic disease (IPA Score: 17), cellular movement (IPA Score: 14) and small molecule biochemistry and metabolism (IPA Score: 12) were constructed, using the list of candidate genes, located in the targeted regions for IBD (Supplementary Table E).

#### Enrichment pathway analysis

The results of the Enrichr analysis showed that several genes within our list appear to be involved in biological processes and/or are molecular components that have been associated or directly/indirectly involved in human IBD. Results of the pathway association analysis using different databases and details of pathways and genes involved in each pathway are shown in Supplementary Table F.

### Association of variants with treatment response

The majority of SNPs identified in the candidate regions for IBD were present in FRD, ARD and the steroid treated group, including SRD and those dogs that were euthanized (NRS/PTS). Several missense and modifier SNPs were present in either FRD and ARD (20 SNPs) or the steroid treated group (10 SNPs) including SRD and NRS/PTS dogs, however, these SNPs were present in only one or two IBD cases (more details are shown in Supplementary Table F).

## Discussion

In the present study, we performed a targeted NGS experiment in previously identified candidate region for canine IBD susceptibility, in order to detect good candidate genes and mutations. The vast majority of variants identified were novel variants. Some of the SNPs identified were not in HWE. Deviation from HWE in case-control genetic association studies is indicative of genetic association (Ziegler et al., 2011). However, intensive selective breeding during breed formation, and therefore loss of random mating that would normally enrich gene pool and maintain HWE, may also explain why some of the SNPs were not in HWE (Ziegler et al., 2011). In addition, due to the small sample size used in this study and missing calls in some of the SNPs that could affect the significance of each genotype, some SNPs may not be in HWE. It is worth mentioning that a number of SNPs identified in the case population were only seen in the heterozygote state, such as those in *EEF1A1* (Elongation Factor 1 Alpha). One possible explanation could be that homozygote SNPs might have been present but have not been captured (low quality/ filtered out etc). Another possibility is that the heterozygotes were introduced recently by either random mutation or outbreeding of GSDs, as heterozygosity in each breed is very low (Lindblad-Toh et al., 2005). It may also be possible that a homozygote state could be lethal.

Two novel stop-gained SNPs in *EEF1A*, were identified only in cases and were present in 26 out of 28 cases. EEF1A1 is an important protein that initiates protein translation elongation and triggers the initiation of protein translation elongation (Kapp and Lorsch 2004; Schulz et al., 2014). Apart from its canonical function in translation elongation by ribosomes, EEF1A1 plays an important role in a wide variety of cellular processes including signaling transduction, heat shock response, cytoskeleton regulation (Negrutskii et al., 2012) and cellular apoptosis (Kobayashi and Yonehara 2009). It has been also documented that binding of EEF1A1 to STAT3 is crucial for STAT3 phosphorylation and for NF-κB/STAT3 activation, which enhances IL-6 expression (Schulz et al., 2014). Elevated levels of this cytokine were reported to play a pivotal role in the initiation of inflammatory processes and progression of disease in many clinical conditions including rheumatoid arthritis, Alzheimer’s disease and Crohn’s disease (Ito 2004; Cacquevel et al., 2004; Nishimoto 2006; Murphy et al., 2012). However, no previous association with IBD has been reported.

*MDH2* (Mitochondrial malate dehydrogenase) is another good candidate gene located on Chr 9, since several missense mutations were detected in this gene, and were only present in IBD cases. In addition, IPA analysis showed that this gene was part of a gene network involved in hereditary disorders and metabolic diseases, as well as involvement in pathways associated with metabolism. The mitochondrial malate dehydrogenase, encoded by *MDH2*, is a mitochondrial protein, which plays an important role in energy production. Altered expression of MDH2 has been reported in studies investigating differentially expressed proteins in intestinal biopsies of IBD patients. Down-regulation of mitochondrial proteins involved in energy production including MDH2 in the colonic mucosal biopsies of ulcerative colitis (UC) patients was previously reported by Hsieh et al. (Hsieh et al., 2006). Results of this study suggested the implications of colonocyte mitochondrial dysfunction and perturbed mucosa immune regulation in the pathogenesis of UC.

In the control population, two known SNPs (rs850782880 and rs852254668) that were identified within less than 300bp downstream of the *MRPS2* gene, were of particular interest. These two SNPs were present in all 20 control GSDs used in this study, therefore could potentially be considered as protective variants for IBD in GSDs. *MRPS2* encodes mitochondrial ribosomal protein subunit 2, which is involved in protein synthesis within the mitochondrion. Most of the mitochondrial proteins including the ribosomal proteins and translation factors that are responsible for the expression of the mitochondrial genome, are synthesized on cytoplasmic ribosomes and imported into mitochondria post-translationally. The mitochondrial oxidative phosphorylation system, which produces the bulk of ATP for almost all eukaryotic cells to sustain cells’ normal functions, depends on the translation of 13 mitochondrial DNA (mtDNA)-encoded polypeptides by mitochondria-specific ribosomes in the mitochondrial matrix. All these peptides are members of the oxidative phosphorylation complexes (Ojala et al., 1981). Several genetic mutations in nuclear genes coding for mitochondrial proteins or mitochondrial genes that can cause defects in mitochondrial transcripts or mitochondrial proteins leading to mitochondrial dysfunction and consequently impaired energy production, have been linked to many inherited diseases (reviewed in (Vafai and Mootha 2012)). Mutations affecting *MRPS2* were observed to cause mitochondrial disorder, altered cellular metabolism, developmental delay, and multiple defects in the oxidative phosphorylation system in a study by Gardeitchik and colleagues (Gardeitchik et al., 2018).

Given that most cellular functions as well as tight junction maintenance and maintenance of the epithelial barrier are energy-dependent, it could be assumed that mitochondrial dysfunction may play a key role in both the onset and recurrence of IBD. The intestinal mucosa of IBD patients is in a state of energy deficiency, characterized by alterations in the oxidative metabolism of epithelial cells and reduced levels of ATP within the intestine (Roediger 1980; Kameyama et al., 1984; Fukushima and Fiocchi 2004). Several studies have provided evidence of mitochondrial stress and abnormalities within the intestinal epithelium of patients with IBD and mice with experimentally induced colitis (Delpre et al., 1989; Nazli et al., 2004; Rodenburg et al., 2008). The hallmarks of mitochondrial dysfunction, including oxidative stress and impaired ATP production, have been observed in the intestines of patients with IBD (Kruidenier and Verspaget 2002; Pravda 2005; Rezaie, et al., 2007) however, it is yet unclear whether these processes occur as a cause or consequence of the disease.

On Chr11, we identified several SNPs within 1 Kb up- and downstream of genes. Two SNPs with modifier impact within 1 kb upstream of *IL3*, 2 SNPs downstream of *PDLIM* and *IL13* and one novel SNP downstream of *IL4*, were identified. Interestingly, all these genes have previously been shown to be associated with human IBD (Jostins et al. 2012). According to our results, the same genes identified as potentially good candidates for IBD in GSDs, further supporting the usefulness of the domestic dog as a natural animal model for human diseases and especially for IBD.

Although disease-associated genetic variations are commonly thought to affect the coding regions of genes, it has been observed that some may alter normal gene expression (Kleinjan and van Heyningen 2005). Thus, it might be possible that the identified SNPs may alter the expression of the genes. It is worth noting that altered expression of interleukins at mRNA and protein levels in human and canine IBD has been reported in several studies.

In addition, there is evidence that a conserved noncoding element (CNE) located between *IL4* and *IL13* controls expression of both genes, as well as *IL5*. A conserved element was identified by cross-species sequence comparison in the intergenic space between the *IL4* and *IL13* cytokine genes. Targeted deletion in mice revealed it to be a coordinate enhancer for *IL4* and *IL13,* as well as for the more distant *IL5* gene. This deletion was also affecting the gene expression in Th2 cells (Loots 2000; Mohrsi et al., 2001).

We also assessed whether there is a correlation between variants identified in the case population and response to treatment. The majority of identified SNPs were present in all treatment response groups and therefore we were unable to detect statistically significant differences. Several missense and modifier SNPs were present in either FRD and ARD or SRD and NRS (PTS) group however, these SNPs were present in only one or two cases. Further studies using a bigger sample size are needed to confirm these results, since the lack of association between SNP markers and response to treatment in the present study maybe attributable to small sample size.

Canine IBD, similar to the condition in humans, is considered to be a complex multifactorial disorder that seems to occur in genetically susceptible individuals after exposure to one or more environmental triggers. In general, it is believed that a number of “susceptibility variants” may cause a general predisposition to IBD, and additional genetic variation or environmental factors may influence specific phenotypic characteristics of the individual such as disease site, disease behaviour or response to treatment. In humans, it has been shown that some susceptibility loci are shared between CD and UC, the two major subtypes of IBD, but others are specific to either CD or UC, which perhaps are responsible for diverging disease courses (Satsangi et al., 1997; Ahmad et al., 2001; Abraham and Cho 2009).

The results of the present study suggest that the sample size of 28 IBD cases may not have enough power to detect associations of rare alleles with the disease. A larger genotyped population may be necessary due to the complex genetic architecture of the disease and environmental effects. In addition, studying environmental differences in dogs with different responses to treatment may help to identify environmental factors that could affect response to treatment. In humans, a number of associations have been reported between environmental factors such as infections in childhood, diet and smoking, and increased risk of developing IBD and their effect on the efficacy of treatments (Wurzelmann et al., 1994; Chapman-Kiddell et al., 2010; Sandborn et al., 2010; Kiss et al., 2011; Sandborn et al., 2015). Further investigation of potential environmental associations, such as deworming and dietary history, vaccination, as well as previous occurrences of infection in GSDs might therefore help identifying factors affecting treatment response.

By performing targeted NGS of the two associated regions identified by GWAS (Peiravan et al., 2018), an attempt was made to identify variants contributing to susceptibility or resistance to canine IBD and then evaluate their correlation with response to treatment. Here, a number of good candidate SNPs with strong association and a potential functional effect were identified. While one or several of these variants may be the causal variant(s), it is also possible that actual causal variants may have been missed in the targeted re-sequencing process or in the genotyping process for technical reasons. Considering the limited sample size of this study, missing calls in some of the SNPs could affect the significance of the results. Therefore, it may be useful to genotype the cases and controls that have not been called properly at these positions, before performing further investigations. The actual functional variant may also be an indel or CNV which has not been investigated in the current study.

A follow-up study in a larger population of IBD cases with different treatment responses is essential to validate results and confirm the variants and genes significantly associated with disease. The SNPs detected by NGS could be further genotyped using a custome made genotyping platform such as Sequenom MassARRAY iPLEX in a larger population of GSDs. In addition, targeted NGS of the other two associated regions on Chr 7 and 13, identified by GWAS will help to identify causal variants and subsequent functional analysis of the causal variants may reveal insights into mechanisms involved in the pathogenesis of canine IBD.

The results presented here represent a starting point for further studies of genetic factors associated with canine IBD. Further studies are necessary to conclusively define whether there is a correlation between certain sets of variants, including newly identified variants and previously known variants in Toll-like receptors (TLR)4, 5 and Nucleotide-binding oligomerization domain-containing protein 2 (NOD2) and response to treatment in GSDs with IBD. Identification of variants associated with the disease could potentially lead to the development of a genetic screening test to assist veterinarians with a diagnosis of IBD, and screening for SNPs that are predictive of response to a specific therapy could, potentially maximize treatment efficiency.

Given the heterogeneity of IBD, it is unlikely that any single marker or class of markers could successfully predict response to treatment. However, a combination of several classes of markers, including genetic, serological and inflammatory markers, may have valuable potential to predict the outcome of a treatment.

## Supporting information

Supplementary A1: Scripts and filtering criteria

Results of quantification of gDNA samples

Concentration and size of amplified libraries for each sample

BAMStat reports for the target region on Chr 9 and 11

Variants on chromosome 9

Variants on chromosome 11

IPA analysis results

Results of Enrichr

Association of variants with treatment response

## Aknoledgement

This work was funded through a BBSRC iCASE studentship (BB/J01236X/1) awarded to K.A. and D.W. with Laboklin GmBH (Bad Kissingen, Germany) as industrial partner as well as the American Kennel Club. We are grateful to owners of GSD who gave permission for their dogs to participate in this study.

