## Supplementary A1: Scripts and filtering criteria for "Targeted next-generation sequencing of Candidate Regions Identified by GWAS Revealed SNPs Associated with IBD in GSDs"

#### Calling variants

java -jar GenomeAnalysisTK.jar -T HaplotypeCaller -R reference.fa -V BQSR.bam -o BQSR.g.vcf –ERC GVCF

#### Joint genotyping

java -jar GenomeAnalysisTK.jar –T GenotypeGVCFs -R reference.fa -o Case_raw.g.vcf --dbSNP -V Case1_BQSR.g.vcf –V Case2_BQSR.g.vcf –V Case3_BQSR.g.vcf –V Case4_BQSR.g.vcf –V Case5_BQSR.g.vcf –V Case6_BQSR.g.vcf –V Case7_BQSR.g.vcf.... –V Case28_BQSR.g.vcf

#### Extraction of raw SNPs

java -jar GenomeAnalysisTK.jar -T SelectVariants -R reference.fa -V Case_raw.g.vcf -selectType SNP -o Case_raw_snps.vcf

#### Filtering criteria.

- QualByDepth (QD) < 2.0

java -jar GenomeAnalysisTK.jar -T VariantFilteration -R reference.fa -V Case_raw_SNP.vcf –filterExpression ‘QD<2.0’ –filtername “FAILED” -o Case_filtered_snps_QD.vcf

- FisherStrand (FS) > 60.0

java -jar GenomeAnalysisTK.jar -T VariantFilteration -R reference.fa -V Case_filtered_snps_QD.vcf –filterExpression ‘FS>60.0’ –filtername “FAILED” -o Case_filtered_snps_QD_FS.vcf

- RMSMappingQuality (MQ) < 40.0

java -jar GenomeAnalysisTK.jar -T VariantFilteration -R reference.fa -V Case_filtered_snps_QD_FS.vcf –filterExpression ‘MQ<40.0’ –filtername “FAILED” -o Case_filtered_snps_QD_FS_MQ.vcf

- MappingQualityRankSumTest (MQRankSum) <-12.5

java -jar GenomeAnalysisTK.jar -T VariantFilteration -R reference.fa -V Case_filtered_snps_QD_FS_MQ.vcf –filterExpression ‘MQRS<-12.5’ –filtername “FAILED” -o Case_filtered_snps_QD_FS_MQ_MQRS.vcf

- ReadPosRankSumTest (ReadPosRankSum) < -8.0

java -jar GenomeAnalysisTK.jar -T VariantFilteration -R reference.fa -V Case_filtered_snps_QD_FS_MQ_MQRS.vcf –filterExpression ‘RPRS<-8.0’ –filtername “FAILED” -o Case_filtered_snps_QD_FS_MQ_MQRS_RPRS.vcf

- StrandOddsRatio (SOR) > 3.0

java -jar GenomeAnalysisTK.jar -T VariantFilteration -R reference.fa -V Case_filtered_snps_QD_FS_MQ_MQRS_RPRS.vcf –filterExpression ‘SOR>3.0’ –filtername “FAILED” -o Case_filtered_snps_QD_FS_MQ_MQRS_RPRS_SOR.vcf
