## Supplementary material for "Targeted next-generation sequencing of Candidate Regions Identified by GWAS Revealed SNPs Associated with IBD in GSDs": Results of quantification of gDNA samples

### Supplementary Table A2: Results of quantification of gDNA samples.

Volume needed for sequencing and average shearing size. VOL: volume, Avg: average.

| Cases/ Controls ID | Qubit (ng/uL) | Vol for 250ng | Shear 1 complete? | Avg shear size (bp) |
| --- | --- | --- | --- | --- |
| GSD35D | 66.0 | 3.8 | Yes | 186 |
| GSD36D | 99.2 | 2.5 | Yes | 177 |
| GSD91D | 60.0 | 4.2 | Yes | 192 |
| GSD96D | 56.0 | 4.5 | Yes | 187 |
| GSD101D | 28.8 | 8.7 | Yes | 183 |
| GSD105D | 6.96 | 35.9 | Yes | 177 |
| GSD108D | 71.8 | 3.5 | Yes | 178 |
| GSD110D | 39.0 | 6.4 | Yes | 175 |
| GSD116C | 43.8 | 5.7 | Yes | 166 |
| GSD127C | 53.4 | 4.7 | Yes | 170 |
| GSD130C | 68.2 | 3.7 | Yes | 189 |
| GSD131C | 364.0 | 0.7 | Yes | 181 |
| GSD132C | 292.0 | 0.9 | Yes | 193 |
| GSD135C | 137.0 | 1.8 | Yes | 187 |
| GSD137C | 47.0 | 5.3 | Yes | 183 |
| GSD138C | 57.0 | 4.4 | Yes | 186 |
| GSD139C | 13 | 19.2 | Yes | 171 |
| GSD140C | 52.4 | 4.8 | Yes | 184 |
| GSD141C | 81.2 | 3.1 | Yes | 178 |
| GSD142C | 25 | 10.0 | Yes | 162 |
| GSD50D | 45.0 | 5.6 | Yes | 162 |
| GSD68D | 20.2 | 12.4 | Yes | 180 |
| GSD77D | 13.5 | 18.5 | Yes | 176 |
| GSD79D | 12.7 | 19.7 | Yes | 170 |
| GSD80D | 12.0 | 20.8 | Yes | 164 |
| GSD85D | 32.8 | 7.6 | Yes | 166 |
| GSD89D | 21.2 | 11.8 | Yes | 174 |
| GSD90D | 23.0 | 10.9 | Yes | 166 |
| GSD92D | 20.4 | 12.3 | Yes | 170 |
| GSD93D | 12.0 | 20.8 | Yes | 179 |
| GSD94D | 22.2 | 11.3 | Yes | 195 |
| GSD99D | 44.6 | 5.6 | Yes | 181 |
| GSD102D | 14.2 | 17.6 | Yes | 184 |
| GSD104D | 18.0 | 13.9 | Yes | 169 |
| GSD109D | 11.2 | 22.3 | Yes | 184 |
| GSD105C | 20.2 | 12.4 | Yes | 189 |
| GSD118C | 12.7 | 19.7 | Yes | 185 |
| GSD129C | 22.8 | 11.0 | Yes | 185 |
| GSD133C | 41.4 | 6.0 | Yes | 192 |
| GSD134C | 24.6 | 10.2 | Yes | 191 |
| GSD136C | 35.8 | 7.0 | Yes | 161 |
| GSD143C | 41.6 | 6.0 | Yes | 186 |
| GSD73D | 4.7 | 55.4 | Yes | 172 |
| GSD128C | 20.6 | 12.1 | Yes | 162 |
| GSD95D | 75.8 | 3.3 | Yes | 173 |
| GSD103D | 22.2 | 11.3 | Yes | 169 |
| GSD67D | 28.8 | 8.7 | Yes | 180 |
| GSD97D | 15.3 | 16.3 | Yes | 190 |
