## Supplementary material for "Targeted next-generation sequencing of Candidate Regions Identified by GWAS Revealed SNPs Associated with IBD in GSDs": Concentration and size of amplified libraries for each sample

### Supplementary Table A3: Concentration and size of amplified libraries for each sample.

| Sample ID | Amplified Library Concentration (ng/uL) | Total ng | Final library size (bp) |
| --- | --- | --- | --- |
| GSD77D | 36 | 972 | 276 |
| GSD133C | 90 | 2430 | 272 |
| GSD80D | 38.2 | 1031.4 | 269 |
| GSD102D | 84.2 | 2273.4 | 272 |
| GSD79D | 70.4 | 1900.8 | 274 |
| GSD85D | 64.4 | 1738.8 | 263 |
| GSD89D | 63 | 1701 | 267 |
| GSD128C | 66.8 | 1803.6 | 273 |
| GSD118C | 55.2 | 1490.4 | 281 |
| GSD93D | 41.6 | 1123.2 | 274 |
| GSD92D | 44.2 | 1193.4 | 267 |
| GSD129C | 97 | 2619 | 276 |
| GSD50D | 73.6 | 1987.2 | 280 |
| GSD131C | 49.8 | 1344.6 | 272 |
| GSD94D | 63.6 | 1717.2 | 271 |
| GSD68D | 74.2 | 2003.4 | 277 |
| GSD99D | 52.6 | 1420.2 | 267 |
| GSD143C | 44.2 | 1193.4 | 271 |
| GSD136C | 27.4 | 739.8 | 262 |
| GSD67D | 108 | 2916 | 282 |
| GSD109D | 89.2 | 2408.4 | 271 |
| GSD134C | 92.4 | 2494.8 | 273 |
| GSD105C | 71.2 | 1922.4 | 278 |
| GSD103D | 42 | 1134 | 277 |
| GSD97D | 97 | 2619 | 285 |
| GSD95D | 27.6 | 745.2 | 278 |
| GSD35D | 84.8 | 2289.6 | 287 |
| GSD73D | 15.3 | 413.1 | 272 |
| GSD90D | 56.4 | 1522.8 | 264 |
| GSD36D | 83.6 | 2257.2 | 269 |
| GSD101D | 104 | 2808 | 278 |
| GSD96D | 71.4 | 1927.8 | 285 |
| GSD91D | 82 | 2214 | 280 |
| GSD108D | 102 | 2754 | 273 |
| GSD105D | 110 | 2970 | 276 |
| GSD116D | 80.2 | 2165.4 | 272 |
| GSD130C | 82 | 2214 | 270 |
| GSD110D | 110 | 2970 | 269 |
| GSD127C | 34.6 | 934.2 | 255 |
| GSD141C | 46.6 | 1258.2 | 265 |
| GSD142C | 100 | 2700 | 270 |
| GSD140C | 92.2 | 2489.4 | 268 |
| GSD139C | 82.8 | 2235.6 | 271 |
| GSD104D | 90 | 2430 | 269 |
| GSD138C | 46.6 | 1258.2 | 272 |
| GSD132C | 16.6 | 448.2 | 263 |
| GSD135C | 16 | 432 | 256 |
| GSD137C | 77.4 | 2089.8 | 274 |
