## Supplementary material for "Targeted next-generation sequencing of Candidate Regions Identified by GWAS Revealed SNPs Associated with IBD in GSDs": BAMStat reports for the target region on Chr 9 and 11

### Supplementary Table B: BAMStat reports for the target region on Chr 9 and 11

Mapped-reads coverage report for target region on Chr 9 (Cases).

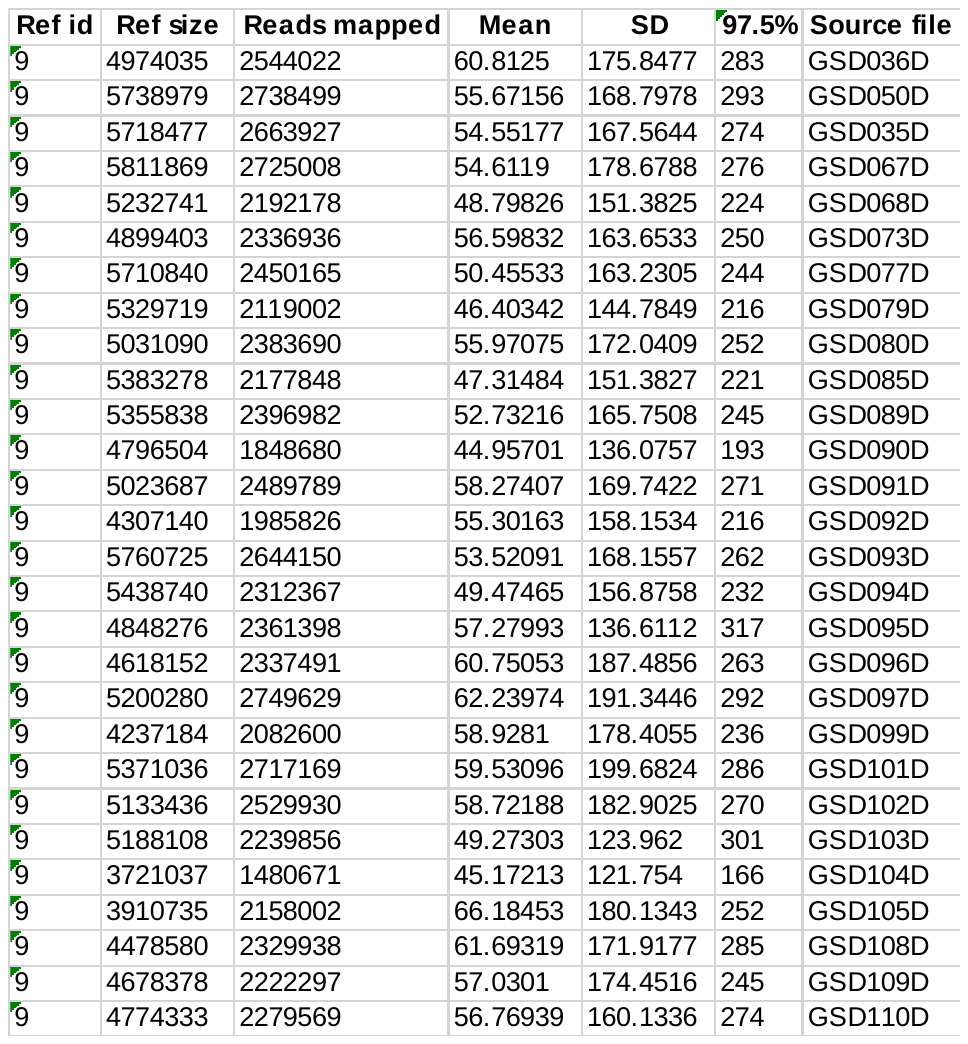

Mapped-reads coverage report for target region on Chr 9 (Controls).

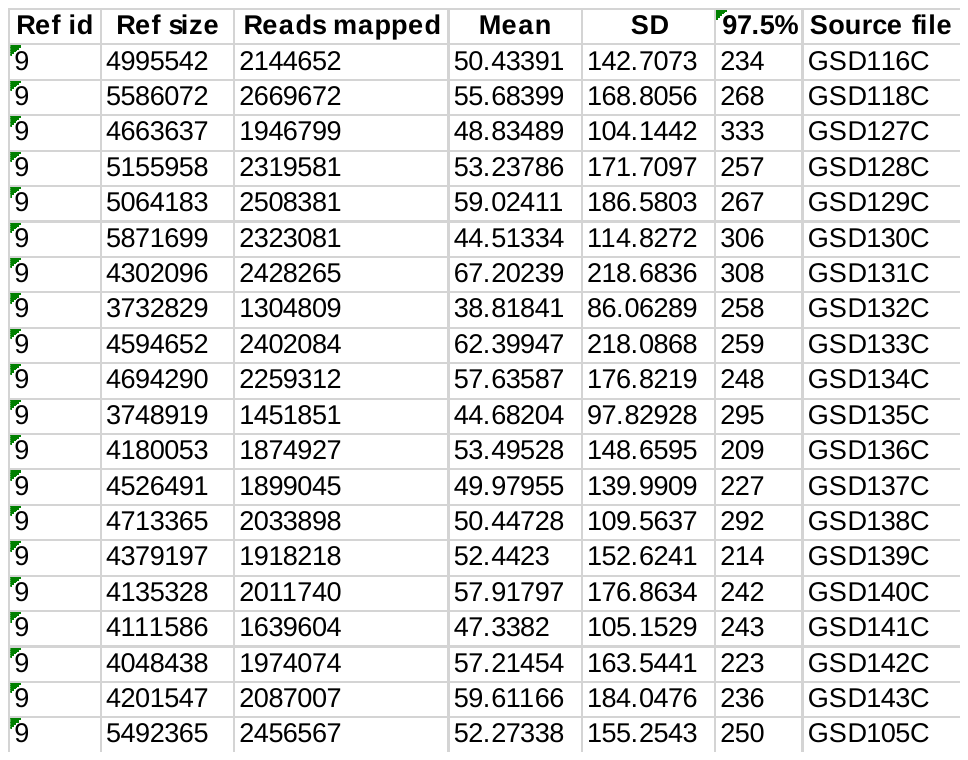

Mapping report for target region on Chr 9 (Cases).

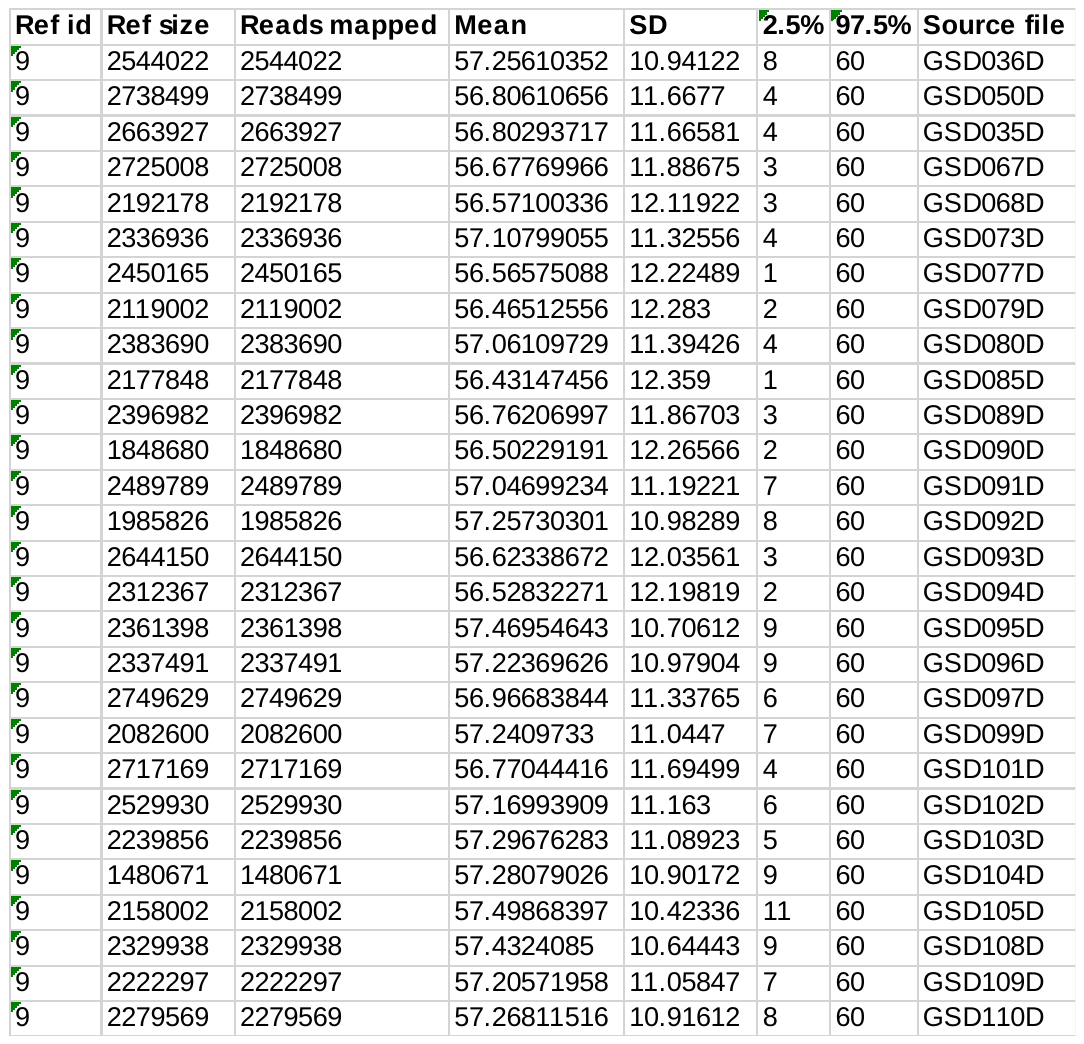

Mapping report for target region on Chr 9 (Controls).

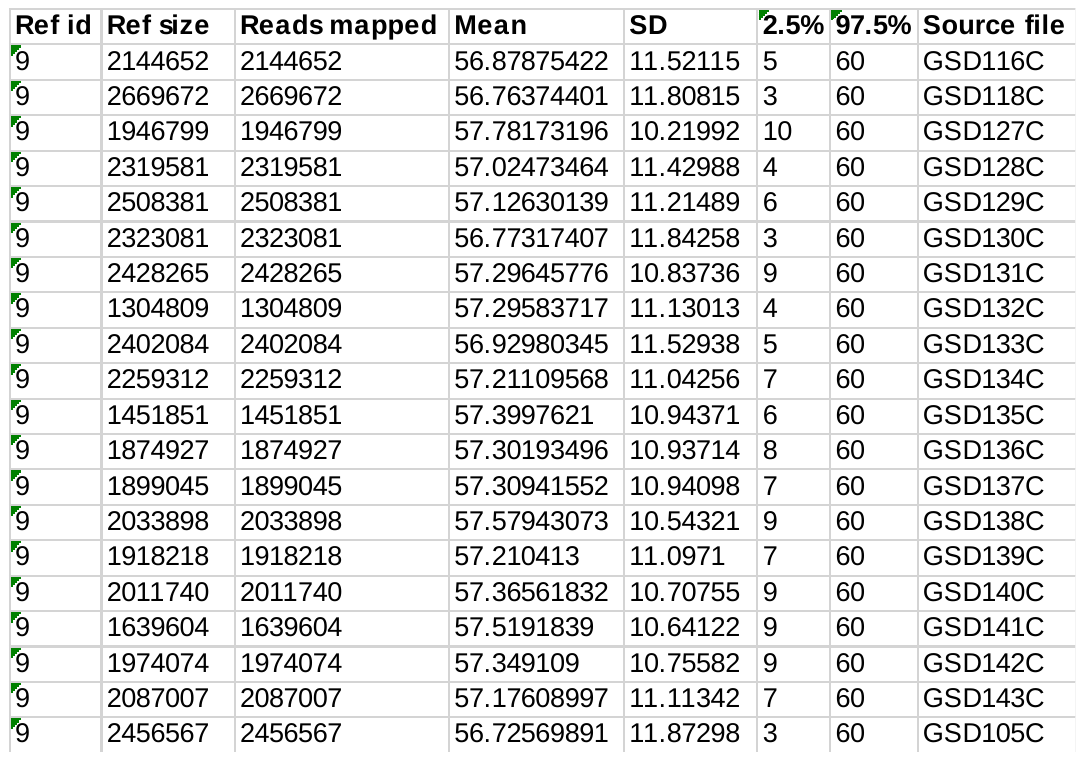

Mapped-reads coverage report for target region on Chr 11 (Cases).

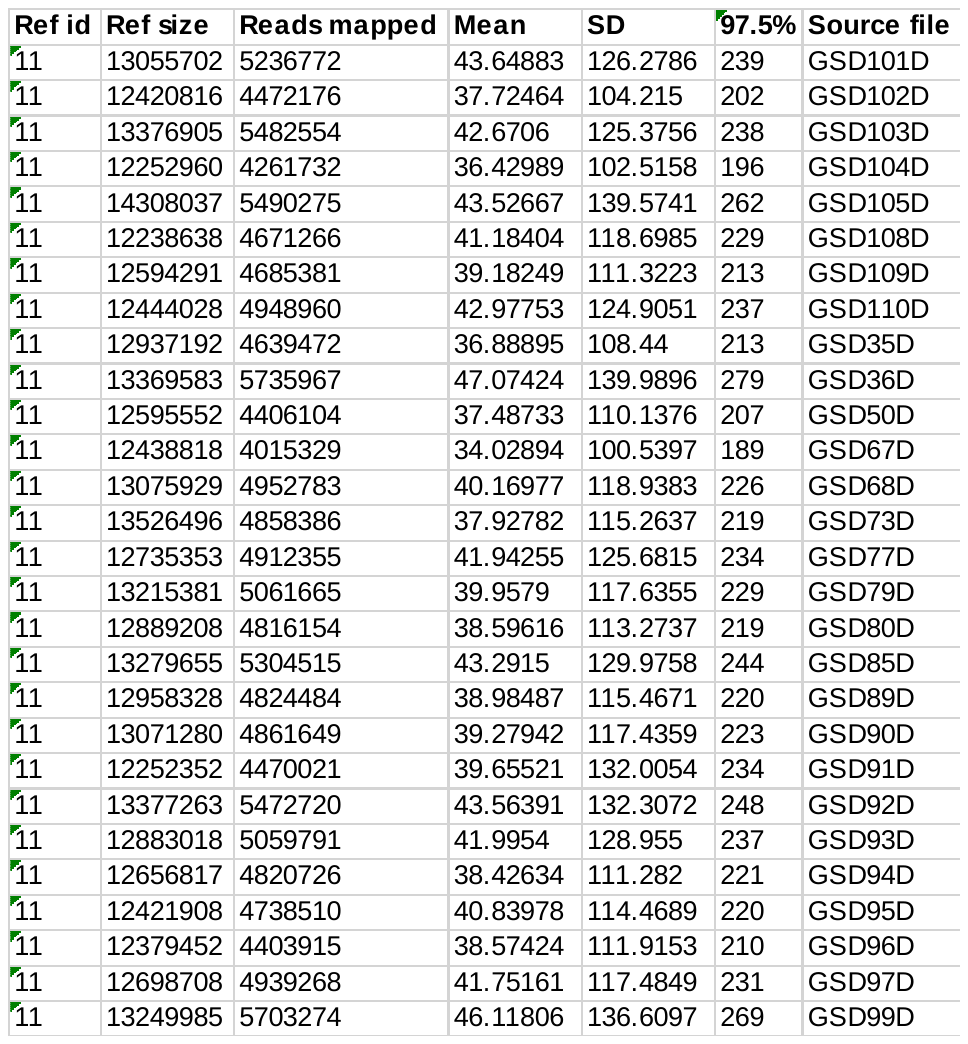

Mapped-reads coverage report for target region on Chr 11 (Controls).

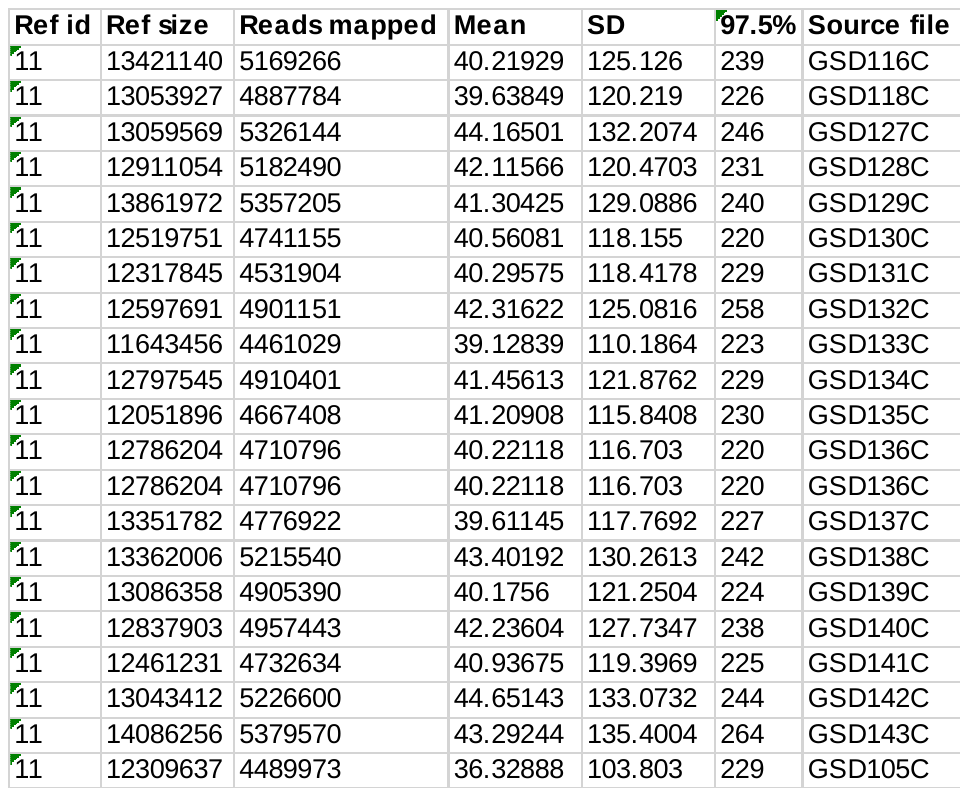

Mapping report for target region on Chr 11 (Cases).

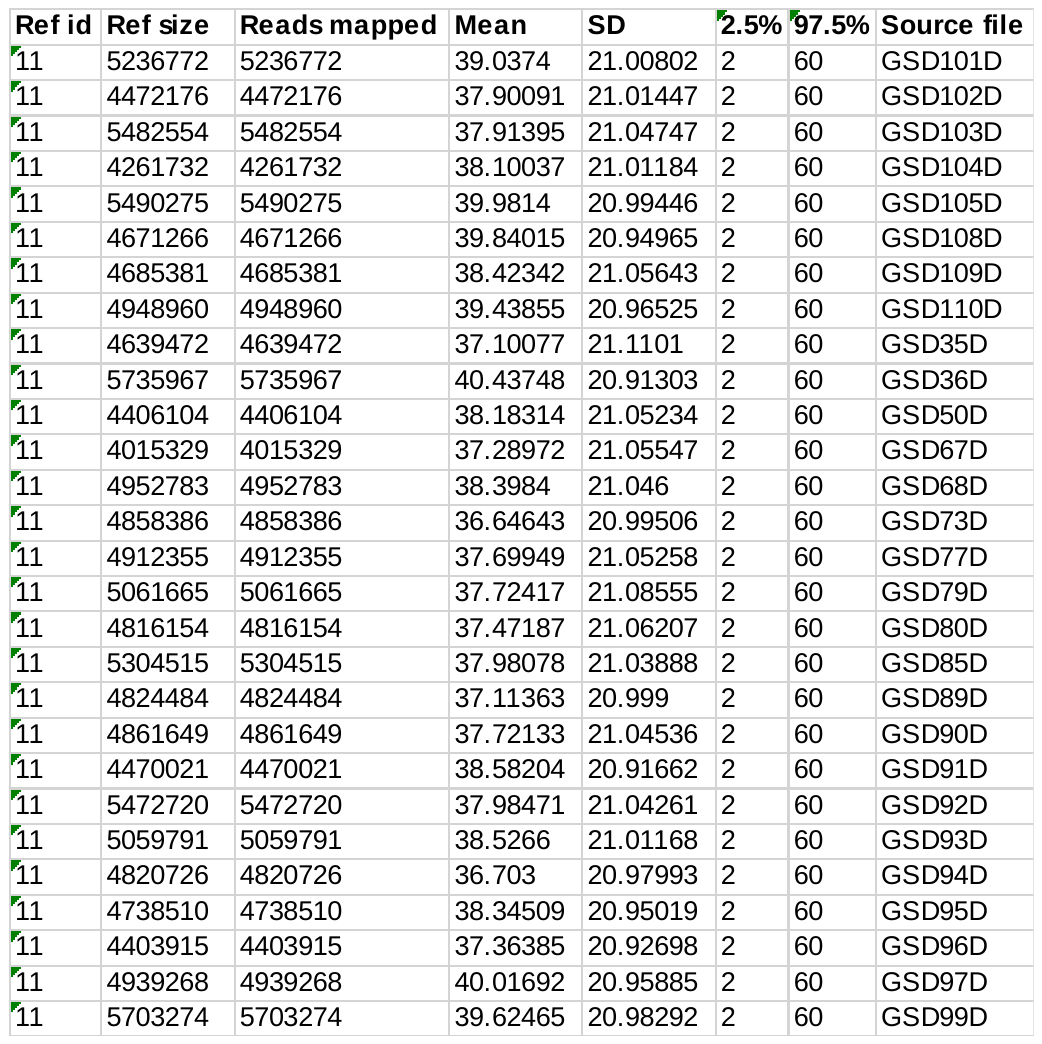

Mapping report for target region on Chr 11 (Controls).

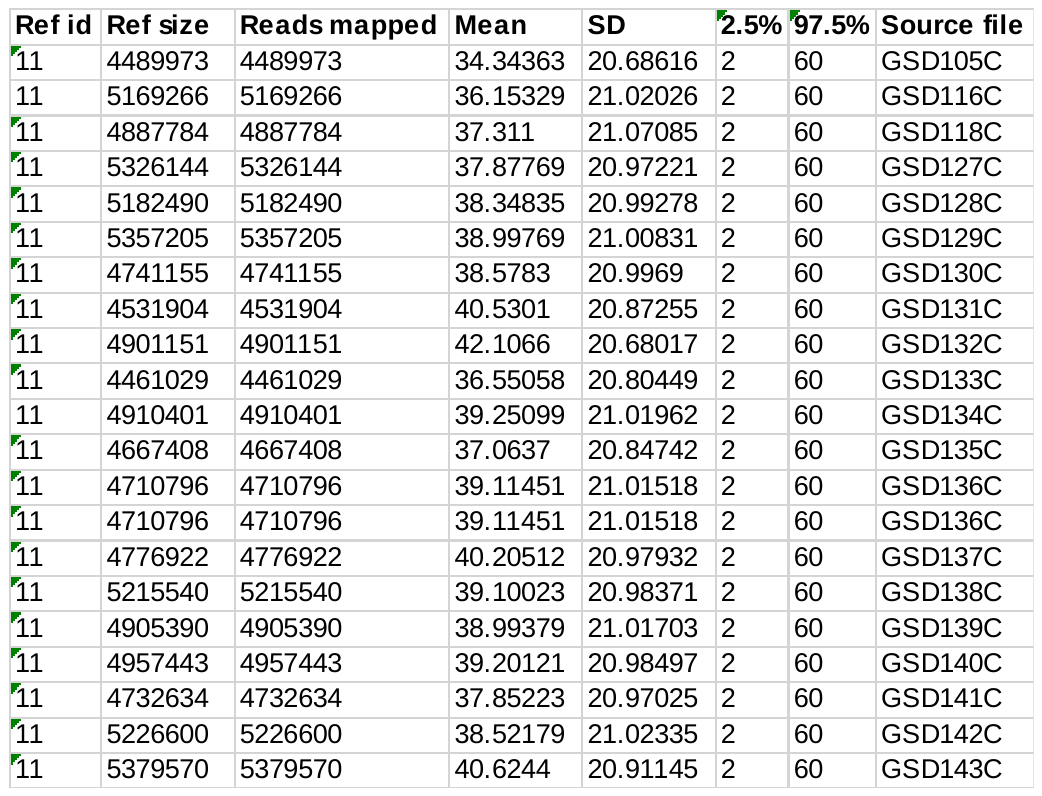
