## Supplementary material for "Targeted next-generation sequencing of Candidate Regions Identified by GWAS Revealed SNPs Associated with IBD in GSDs": Variants on chromosome 9

### Supplementary Table C: Variants on chromosome 9

Variants within genes (high and moderated impact) and within 1 Kb up- and downstream gene (modifier impact) in the case population.

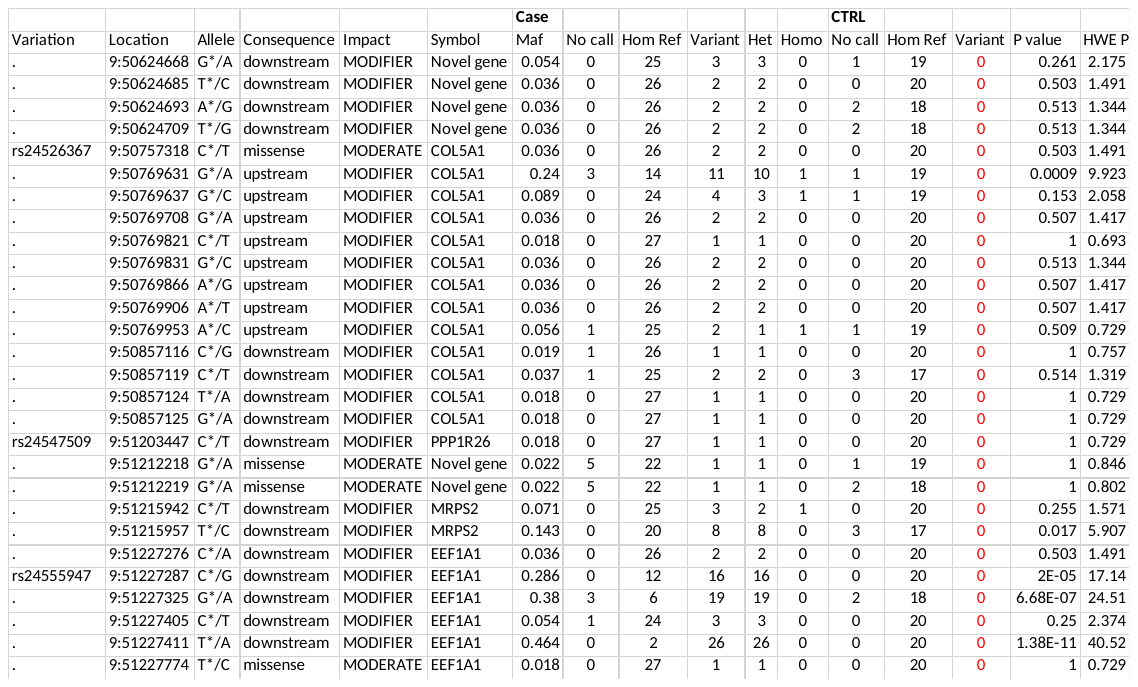

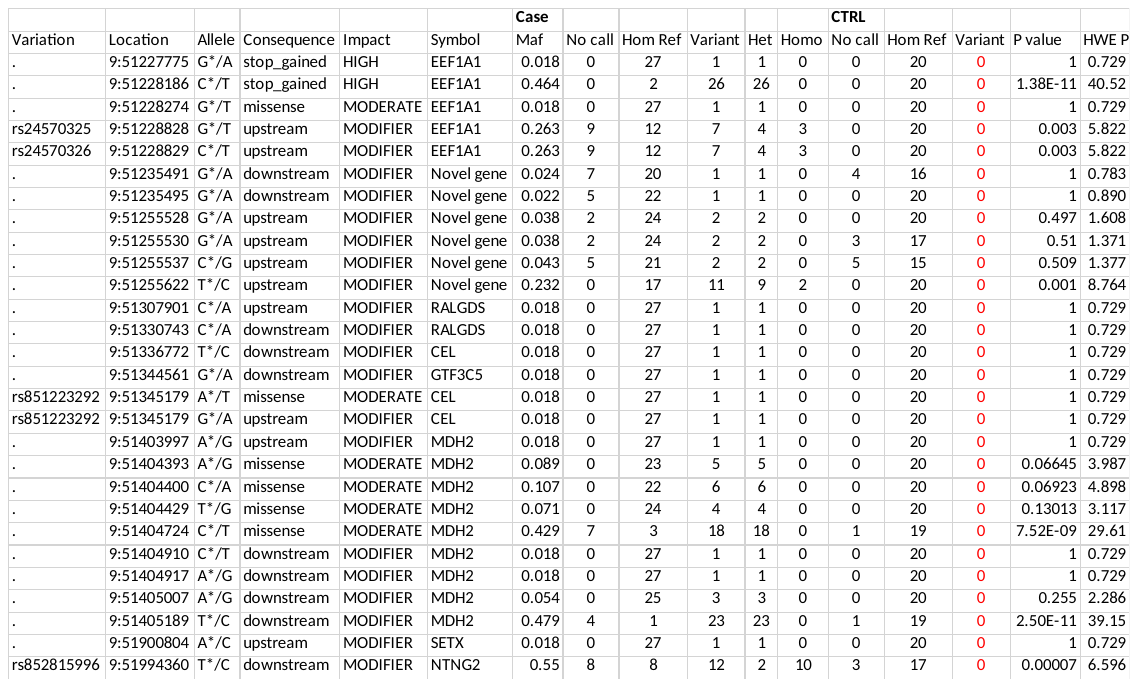

Hom Ref: sites with reference allele (AA), Het: Heterozygous (AB), Homo: Homozygous (BB), P value: Fisher’s exact probability test two tailed p value, Hardy-Weinberg equilibrium: HWE, HWE P: chi-square probability test p value used to test for HWE.

Variants within genes (high and moderated impact) and within 1 Kb up- and downstream gene (modifier impact) in the control population.

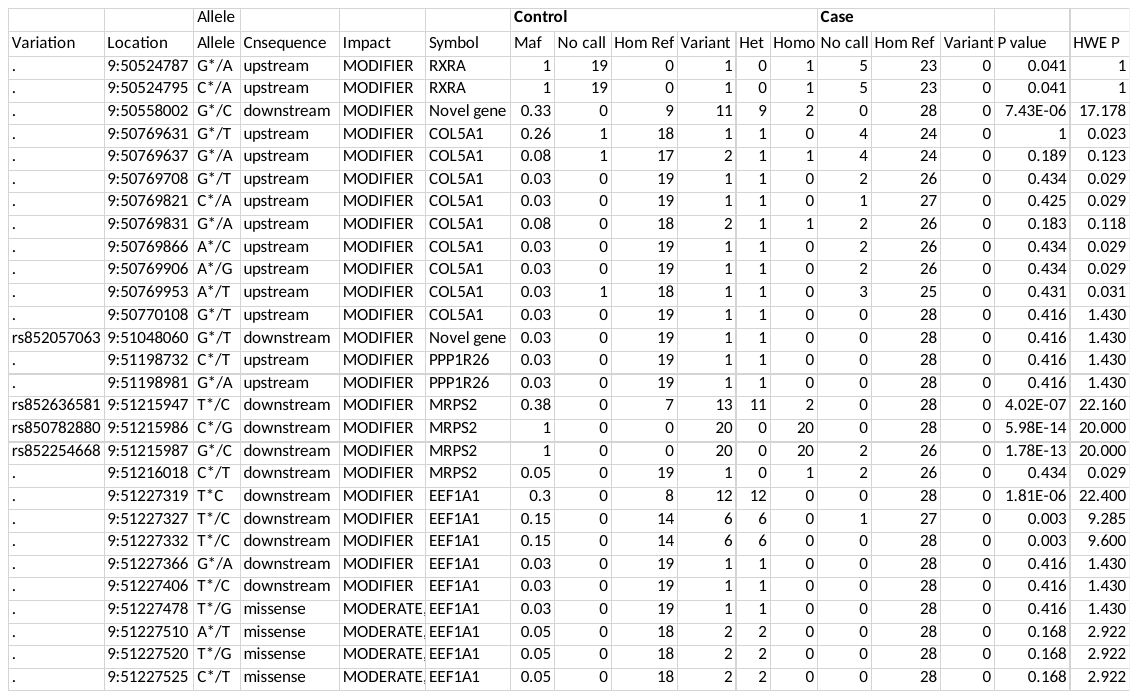

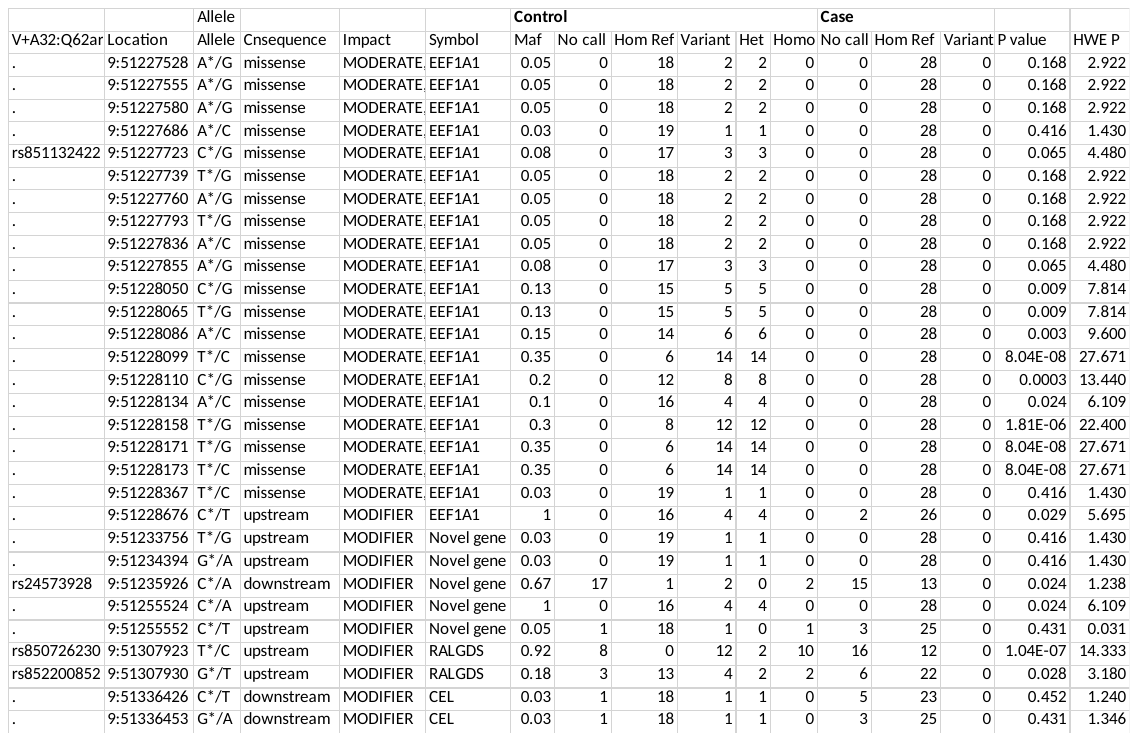

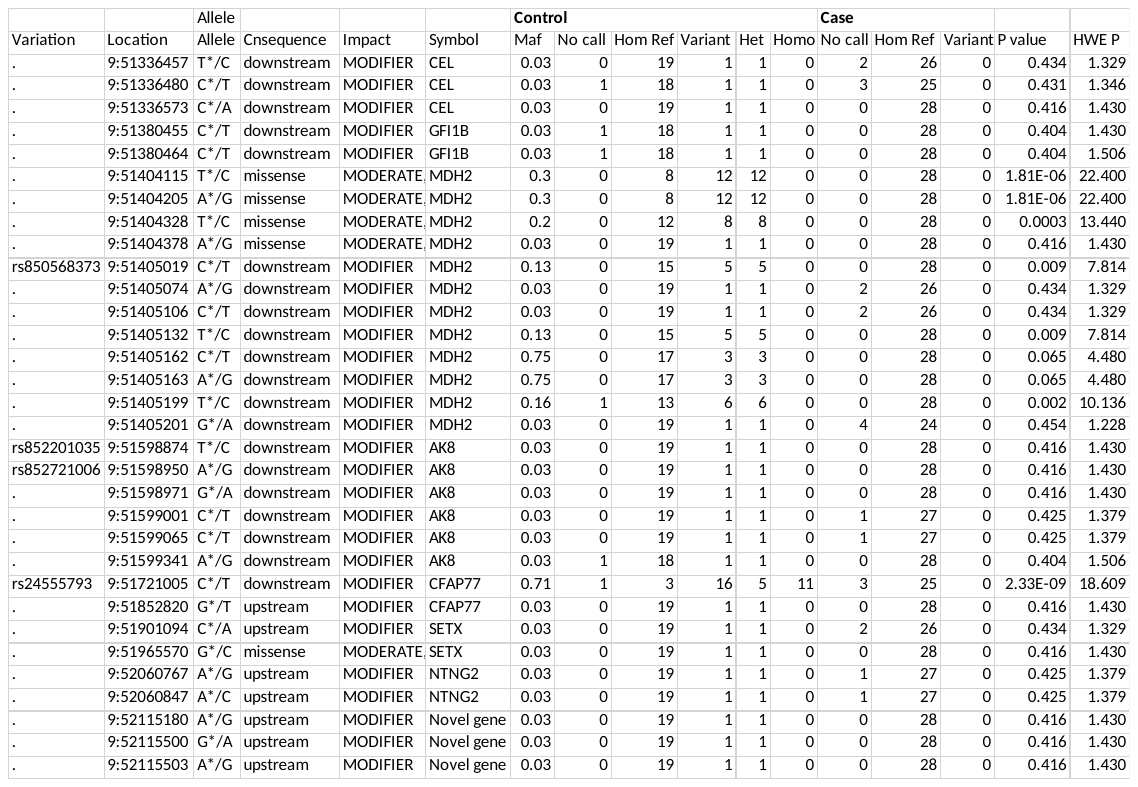

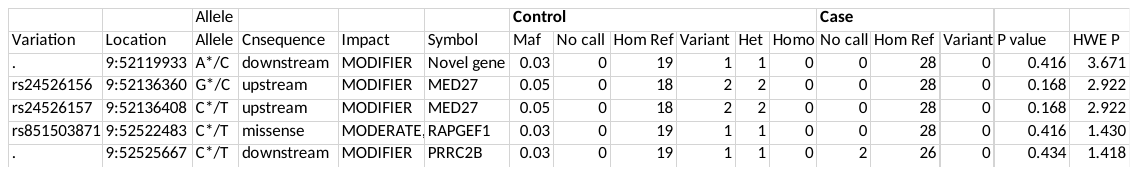

Hom Ref: sites with reference allele (AA), Het: Heterozygous (AB), Homo: Homozygous (BB), P value: Fisher’s exact probability test two tailed p value, Hardy-Weinberg equilibrium: HWE, HWE P: chi-square probability test p value used to test for HWE.

Variants overlapped between the case and the control populations with the same alternate in the both populations.

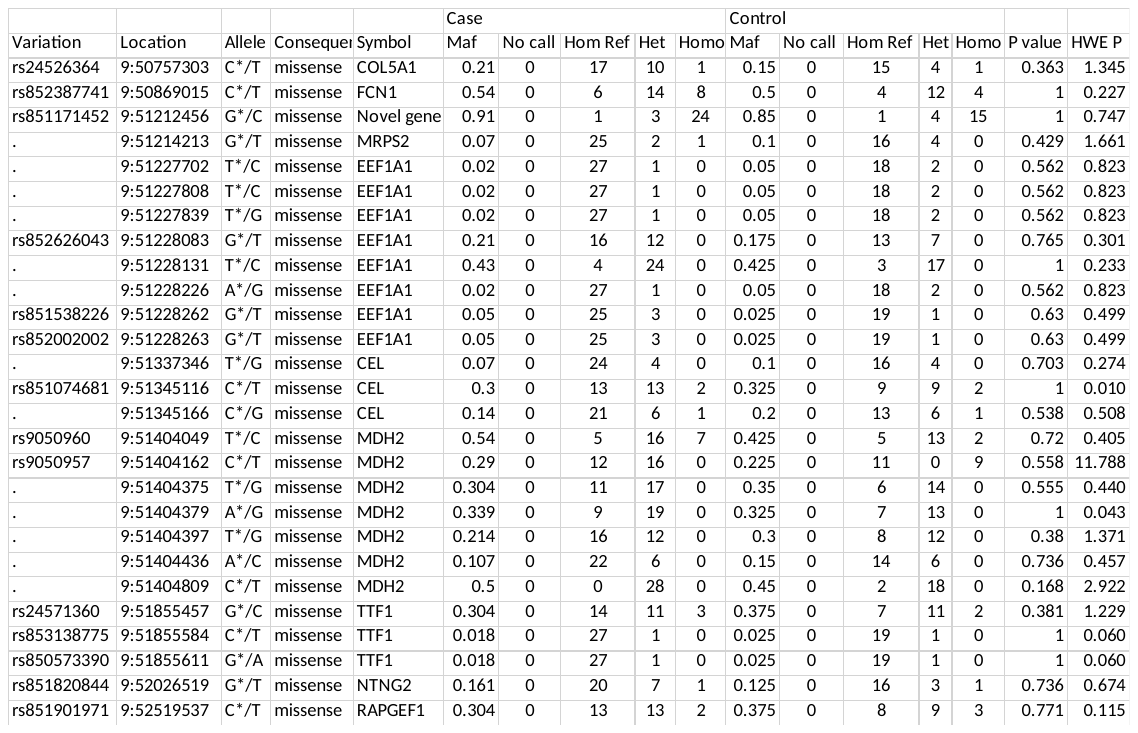

Hom Ref: sites with reference allele (AA), Het: Heterozygous (AB), Homo: Homozygous (BB), P value: Fisher’s exact probability test two tailed p value, Hardy-Weinberg equilibrium: HWE, HWE P: chi-square probability test p value used to test for HWE.
