## Supplementary material for "Targeted next-generation sequencing of Candidate Regions Identified by GWAS Revealed SNPs Associated with IBD in GSDs": Variants on chromosome 11

### Supplementary Table D: Variants on chromosome 11

Variants within genes (high and moderated impact) and within 1 Kb up- and downstream gene (modifier impact) in the case population.

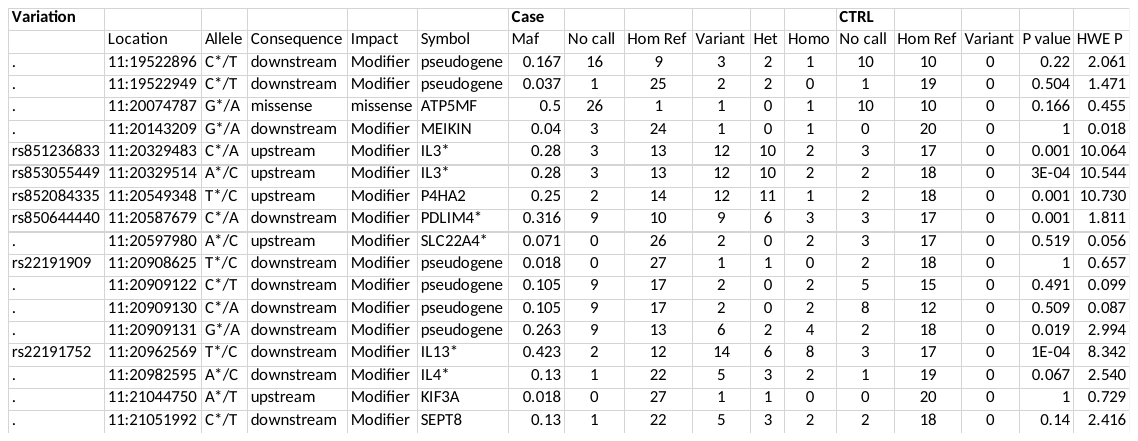
*Genes already found to be associated with human IBD.

Hom Ref: sites with reference allele (AA), Het: Heterozygous (AB), Homo: Homozygous (BB), P value: Fisher’s exact probability test two tailed p value, Hardy-Weinberg equilibrium: HWE, HWE P: chi-square probability test p value used to test for HWE.

*IL3*: Interleukin 3, *P4HA2*: Prolyl 4-hydroxylase subunit alpha-2, *SLC22A4*: Solute Carrier Family 22 Member 4, *KIF3A*: Kinesin Family Member 3A, *MEIKIN*: Meiotic Kinetochore Factor, *PDLIM*: PDZ and LIM Domain, *IL13*: Interleukin 13, *IL4*: Interleukin 4, *SEPT8*: Septin8. *SLC22A4*: Solute Carrier Family 22 Member 4, *KIF3A*: Kinesin Family Member 3A, *MEIKIN*: Meiotic Kinetochore Factor, *PDLIM*: PDZ and LIM Domain, *IL13*: Interleukin 13, *IL4*: Interleukin 4, *SEPT8*: Septin8.

Variants within genes (high and moderated impact) and within 1 Kb up- and downstream gene (modifier impact) in the control population.

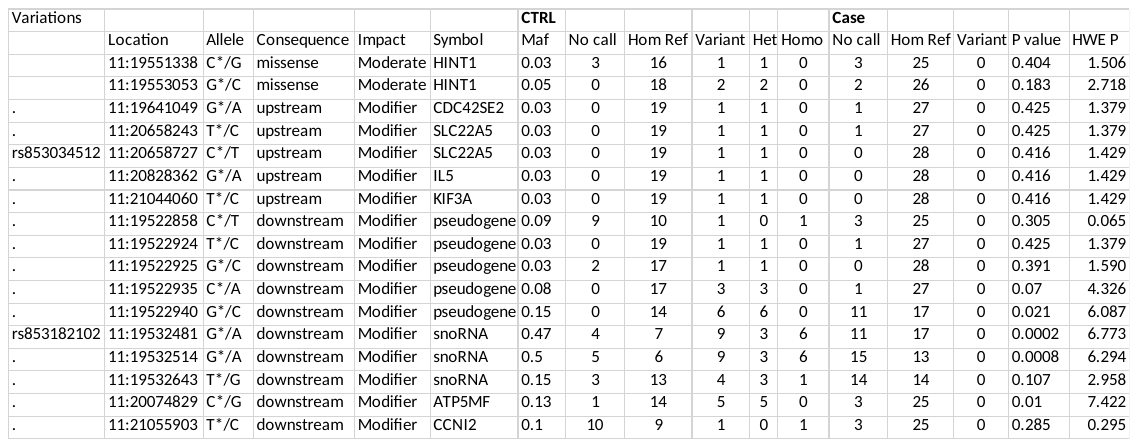

Hom Ref: sites with reference allele (AA), Het: Heterozygous (AB), Homo: Homozygous (BB), P value: Fisher’s exact probability test two tailed p value, Hardy-Weinberg equilibrium: HWE, HWE P: chi-square probability test p value used to test for HWE.

*HINT1:* Histidine triad nucleotide-binding protein 1, *CDC42S2E*: Cell division control protein 42 homolog small effector protein 2, *SLC22A4*: Solute Carrier Family 22 Member 5, *IL5*: Interleukine 5, *KIF3A*: Kinesin Family Member 3A, snoRNA: Small nucleolar RNA, *ATP5MF*: ATP synthase subunit f (Mitochondrial membrane ATP synthase).
