## Supplementary material for "Targeted next-generation sequencing of Candidate Regions Identified by GWAS Revealed SNPs Associated with IBD in GSDs": IPA analysis results

### Supplementary Table E: IPA analysis results.

##### Pathways and genes involved in each pathway.

| Pathway | Genes | Gene description |
| --- | --- | --- |
| Airway inflammation in Asthma | IL4 | interleukin 4 |
|  | IL5 | interleukin 5 |
|  | IL13 | interleukin 13 |
| Haematopoiesis from multipotent stem cells | IL3 | interleukin 3 |
|  | IL4 | interleukin 4 |
|  | IL5 | interleukin 5 |
| Role of cytokines in mediating communication between immune cells | IL3 | interleukin 3 |
|  | IL4 | interleukin 4 |
|  | IL5 | interleukin 13 |
|  | IL3 | interleukin 3 |
| Hepatic cholestasis | IL3 | interleukin 3 |
|  | IL4 | interleukin 4 |
|  | IL5 | interleukin 5 |
|  | IL13 | interleukin 13 |
| Fc Epsilon RI signalling | IL3 | interleukin 3 |
|  | IL4 | interleukin 4 |
|  | IL5 | interleukin 5 |
|  | IL13 | interleukin 13 |
| Role of PRRs in recognition of bacterial and viruses | IL3 | interleukin 3 |
|  | IL4 | interleukin 4 |
|  | IL5 | interleukin 5 |
|  | IL13 | interleukin 13 |
| Haematopoiesis from multipotent stem cells | IL3 | interleukin 3 |
|  | IL4 | interleukin 4 |
|  | IL5 | interleukin 5 |
|  | IL13 | interleukin 13 |
| HMGB1 signalling | IL3 | interleukin 3 |
|  | IL4 | interleukin 4 |
|  | IL5 | interleukin 5 |
|  | IL13 | interleukin 13 |
|  | IL3 | interleukin 3 |
| TH2 pathway | IL3 | interleukin 3 |
|  | IL4 | interleukin 4 |
|  | IL5 | interleukin 5 |
|  | IL13 | interleukin 13 |
| Th1 and Th2 activation pathway | IL3 | interleukin 3 |
|  | IL4 | interleukin 4 |
|  | IL5 | interleukin 5 |
|  | IL13 | interleukin 13 |
| T helper cell differentiation | IL4 | interleukin 4 |
|  | IL5 | interleukin 5 |
|  | IL13 | interleukin 13 |
| Differential regulation of cytokine production in Macrophages and T helper cells by IL_17A and IL_17F | IL3 | interleukin 3 |
|  | IL13 | interleukin 13 |

| **Communication between innate and adaptive immune cells** | IL3 | interleukin 3 |
| --- | --- | --- |
|  | IL4 | interleukin 4 |
|  | IL5 | interleukin 5 |
| **Differential regulation of cytokine production in intestinal epithelial cells by IL_17A and IL_17F** | IL13 | interleukin 13 |
|  | IL3 | interleukin 3 |
| **Glucocorticoid Receptor Signalling** | IL3 | interleukin 3 |
|  | IL4 | interleukin 4 |
|  | IL5 | interleukin 5 |
|  | IL13 | interleukin 13 |
| **autoimmune thyroid disease signalling** | IL4 | interleukin 4 |
|  | IL5 | interleukin 5 |
| **Allograft rejection signalling** | IL4 | interleukin 4 |
|  | IL5 | interleukin 5 |
| **IL3 signalling** | IL3 | interleukin 3 |
|  | RAPGEF1 | rap guanine nucleotide |
| **Crosstalk between dendritic and natural killer cells** | IL3 | interleukin 3 |
|  | IL4 | interleukin 4 |
| **Aspartate degradation II** | MDH2* | aspartate degradation II |
| **Assembly of RNA polymerase III complex** | GTF3C5 | assembly of RNA polymerase III complex |
| **IL12 signalling and production in Macrophages** | IL4 | interleukin 4 |
|  | RXRA* |  |
| **Hepatic fibrosis/hepatic stellate cell activation** | COL5A1* | collagen type V alpha1 |
|  | IL4 | interleukin 4 |
| **Pyrimidine deoxyribonucleotides De Novo biosynthesis** | AK8 | adenylate kinase8 |
| **TCA cycle II (Eukaryotic)** | MDH2* | malate dehydrogenase |
| **Gluconeogenesis I** | MDH2* | malate dehydrogenase |

##### Networks

Hereditary disorder and metabolic disease networks and genes involved in this network. Grey filled shapes represent genes included in the list of candidate genes identified in the targeted regions. Solid and dotted lines represent direct and indirect interaction between genes respectively.

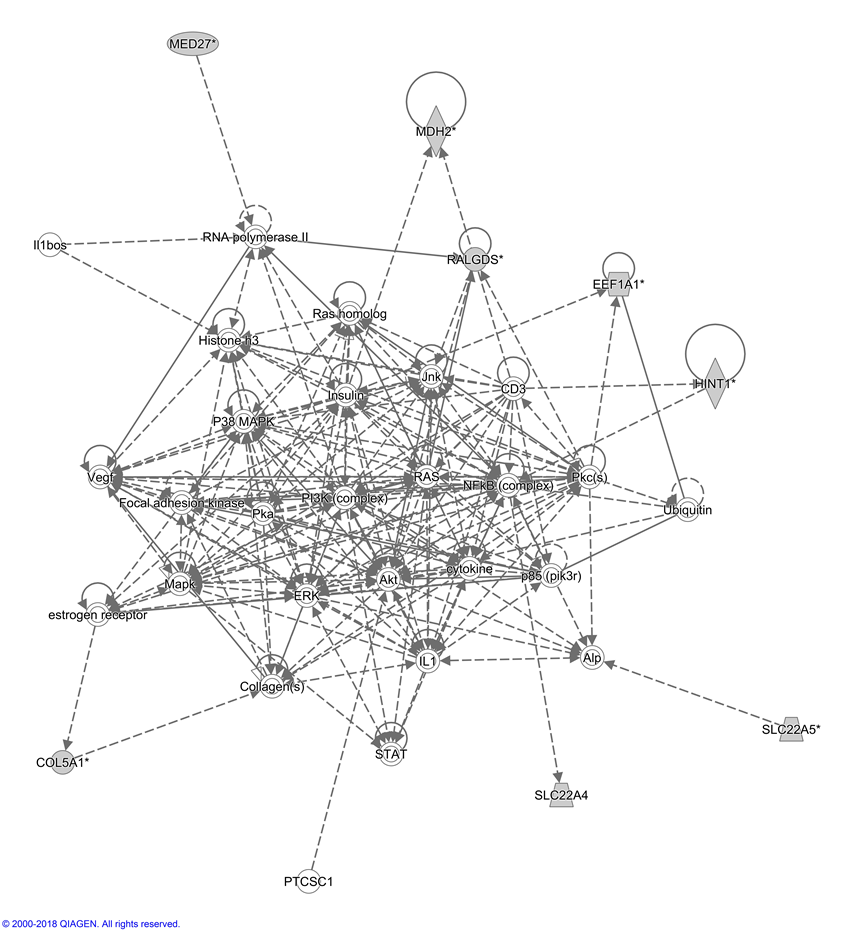

Cellular movement and genes involved in this network.

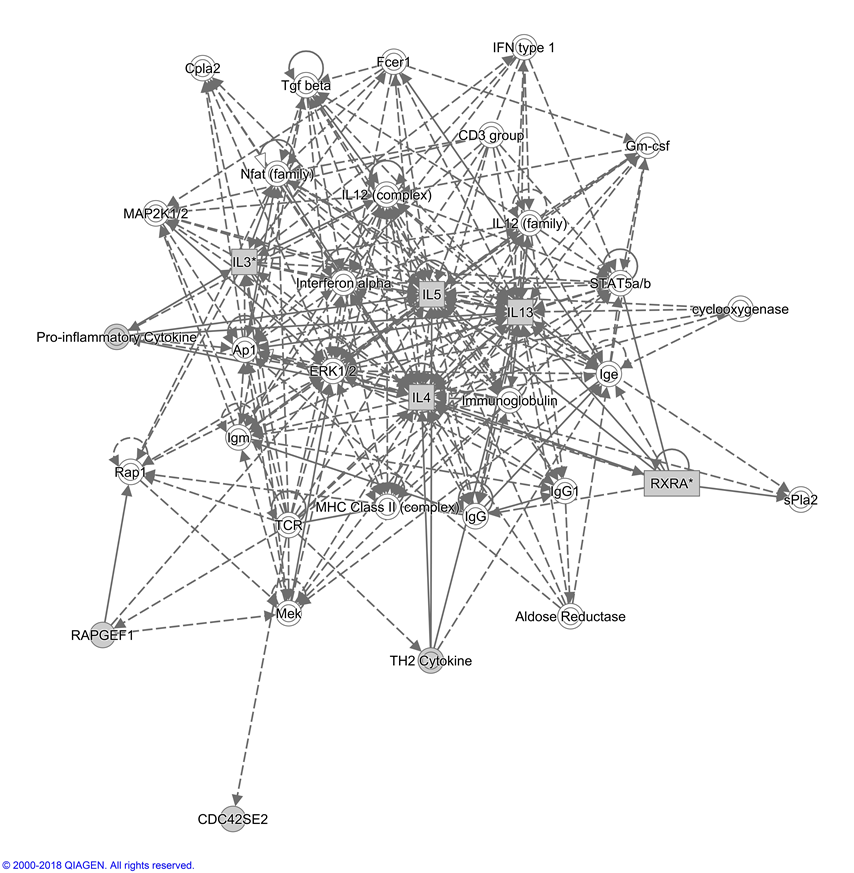
