## Supplementary material for "Targeted next-generation sequencing of Candidate Regions Identified by GWAS Revealed SNPs Associated with IBD in GSDs": Results of Enrichr

**Supplementary Table F: Results of Enrichr.**

| Database | Pathway | P- value | Adjusted | Z score | Genes involved in pathway |
| --- | --- | --- | --- | --- | --- |
| KEGG | Asthma_ airway inflammation | 1.87E-07 | 0.00001011 | -1.63 | IL4;IL3;IL5;IL13 |
|  | Fc epsilon RI signalling pathway | 0.000004642 | 0.0001253 | -2.02 | IL4;IL3;IL5;IL13 |
|  | Jak-STAT signalling pathway | 0.0001283 | 0.002251 | -1.88 | IL4;IL3;IL5;IL13 |
|  | Inflammatory bowel disease (IBD) | 0.0001667 | 0.002251 | -1.86 | IL4;IL5;IL13 |
|  | Choline metabolism in cancer | 0.0006109 | 0.005498 | -1.8 | SLC22A4;SLC22A5;RALGDS |
|  | Hematopoietic cell lineage | 0.0004082 | 0.004408 | -1.67 | IL4;IL3;IL5 |
|  | Cytokine-cytokine receptor interaction | 0.0009112 | 0.007029 | -1.73 | IL4;IL3;IL5;IL13 |
|  | Allograft rejection_ | 0.001788 | 0.01207 | -1.46 | IL4;IL5 |
|  | Intestinal immune network for IgAproduction | 0.00284 | 0.01704 | -1.5 | IL4;IL5 |
|  | Autoimmune thyroid disease | 0.003452 | 0.01864 | -1.51 | IL4;IL5 |
|  | PI3K-Akt signalling pathway | 0.01837 | 0.0763 | -1.76 | IL4;IL3;RXRA |
|  | Rap1 signalling pathway | 0.04723 | 0.1244 | -1.48 | RAPGEF1;RALGDS |
|  | Metabolic pathways | 0.0507 | 0.1244 | -1.49 | MDH2;P4HA2;CEL;AK8;GBGT1 |
| Wikipathways | Cytokines and Inflammatory Response | 8.95E-08 | 0.000005369 | -2.24 | IL4;IL3;IL5;IL13 |
|  | Allograft Rejection | 0.000008037 | 0.0001837 | -2 | IL4;IL5;COL5A1;IL13 |
|  | Cytokines and Inflammatory Response | 0.000009183 | 0.0001837 | -1.92 | IL4;IL5;IL13 |
|  | Inflammatory Response Pathway | 0.001116 | 0.01339 | -1.91 | IL4;IL5 |
|  | IL-3 signalling Pathway | 0.002958 | 0.02535 | -2.09 | IL3;RAPGEF1 |
|  | Hematopoietic Stem Cell Differentiation | 0.002724 | 0.02535 | -1.8 | IL3;IL5 |
|  | IL-5 signalling Pathway | 0.005619 | 0.04214 | -1.88 | IL5;RAPGEF1 |
| Panther2016 | Interleukin signaling pathway | 0.0003815 | 0.002671 | -1.61 | IL4;IL5;IL13 |
|  | Integrin signalling pathway | 0.02725 | 0.05127 | -1.59 | COL5A1;RAPGEF1 |
|  | Vitamin D metabolism and pathway | 0.01313 | 0.04594 | -0.58 | RXRA |
|  | Axon guidance mediated by netrin | 0.04837 | 0.05643 | -0.74 | NTNG2 |
|  | De novo purine biosynthesis | 0.04205 | 0.05643 | -0.64 | AK8 |
