## Supplementary material for "Targeted next-generation sequencing of Candidate Regions Identified by GWAS Revealed SNPs Associated with IBD in GSDs": Association of variants with treatment response

**Supplementary Table G: Association of variants with treatment response**

Variants identified on Chr 9 and cases carrying the variants.

FRD: Food responsive, ARD: Antibiotic responsive, SRD: Steroid responsive, NRS: No response to steroid, PTS: Put to sleep.

| Variation | Location | Consequence | Symbol | Maf | No call | Variant | Het | Homo |
| --- | --- | --- | --- | --- | --- | --- | --- | --- |
| . | 9:50624668 | downstream | Novel gene | 0.054 | 0 | 3 | GSD36D(SRD),GSD91D(SRD),GSD109D(FRD) |  |
| . | 9:50624685 | downstream | Novel gene | 0.036 | 0 | 2 | GSD91D(SRD),GSD109D(FRD) |  |
| . | 9:50624693 | downstream | Novel gene | 0.036 | 0 | 2 | GSD91D(SRD),GSD109D(FRD) |  |
| . | 9:50624709 | downstream | Novel gene | 0.036 | 0 | 2 | GSD91D(SRD),GSD109D(FRD) |  |
| rs24526367 | 9:50757318 | missense | COL5A1 | 0.036 | 0 | 2 | GSD94D(ARD),GSD101D(SRD) |  |
| . | 9:50769631 | upstream | COL5A1 | 0.24 | 3 | 11 | GSD50D(FRD),GSD77D(NRS,PTS),GSD80D(NRS,PTS),  GSD85D(SRD),GSD89D(NRS,PTS),GSD90D(ARD),  GSD91D(SRD),GSD94D(ARD),GSD97D(SRD),GSD102D(SRD) | GSD105D(ARD) |
| . | 9:50769637 | upstream | COL5A1 | 0.089 | 0 | 4 | GSD67D(SRD),GSD85D(SRD),GSD90D(ARD), | GSD95D(FRD) |
| . | 9:50769708 | upstream | COL5A1 | 0.036 | 0 | 2 | GSD79D(FRD),GSD94D(ARD) |  |
| . | 9:50769821 | upstream | COL5A1 | 0.018 | 0 | 1 | GSD105D(ARD) |  |
| . | 9:50769831 | upstream | COL5A1 | 0.036 | 0 | 2 | GSD105D(ARD),GSD110D(SRD) |  |
| . | 9:50769866 | upstream | COL5A1 | 0.036 | 0 | 2 | GSD36D(SRD),GSD95D(FRD) |  |
| . | 9:50769906 | upstream | COL5A1 | 0.036 | 0 | 2 | GSD85D(SRD),GSD95D(FRD) |  |
| . | 9:50769953 | upstream | COL5A1 | 0.056 | 1 | 2 | GSD108D(SRD) | GSD102D(SRD) |
| . | 9:50857116 | downstream | COL5A1 | 0.019 | 1 | 1 | GSD105D(ARD) |  |
| . | 9:50857119 | downstream | COL5A1 | 0.037 | 1 | 2 | GSD73D(FRD),GSD105D(ARD) |  |
| . | 9:50857124 | downstream | COL5A1 | 0.018 | 0 | 2 | GSD73D(FRD),GSD105D(ARD) |  |
| Variation | **Location** | **Consequence** | **Symbol** | **Maf** | **No call** | **Variant** | **Het** | **Homo** |
| . | 9:50857125 | downstream | COL5A1 | 0.018 | 0 | 1 | GSD73D(FRD) |  |
| rs24547509 | 9:51203447 | downstream | PPP1R26 | 0.018 | 0 | 1 | GSD94D(ARD) |  |
| . | 9:51212218 | missense | Novel gene | 0.022 | 5 | 1 | GSD99D(FRD) |  |
| . | 9:51212219 | missense | Novel gene | 0.022 | 5 | 1 | GSD99D(FRD) |  |
| . | 9:51215942 | downstream | MRPS2 | 0.071 | 0 | 3 | GSD73D(FRD),GSD77D(NRS, PTS) | GSD36D(SRD) |
| . | 9:51215957 | downstream | MRPS2 | 0.143 | 0 | 3 | GSD85D(SRD),GSD89D(NRS,PTS),GSD91D(SRD), |  |
| . | 9:51227276 | downstream | EEF1A1 | 0.036 | 0 | 4 | GSD97D(SRD),GSD99D(FRD),GSD109D(FRD),GSD110D(SRD) |  |
| rs24555947 | 9:51227287 | downstream | EEF1A1 | 0.286 | 0 | 16 | GSD35D(FRD),GSD50D(FRD),GSD67D(SRD),GSD79D(FRD),  GSD89D(NRS,PTS),GSD90D(ARD),GSD91D(SRD),GSD93D(SRD),  GSD94D(ARD),GSD95D(FRD),GSD97D(SRD),GSD99D(FRD),  GSD101D(SRD),GSD103D(FRD),GSD104D(ARD),GSD109D(FRD) |  |
| . | 9:51227325 | downstream | EEF1A1 | 0.38 | 3 | 19 | GSD35D(FRD),GSDGSD36D(SRD),GSD50D(FRD),GSD67D(SRD),  GSD68D(NRS,PTS),GSD73D(FRD),GSD79D(FRD),GSD80D(NRS,PTS),  GSD90(ARD),GSD92D(FRD),GSD93(SRD),GSD95D(FRD),  GSD96D(SRD),GSD99D(FRD),GSD101D(SRD),GSD102D(SRD),  GSD103D(FRD), GSD104D(ARD),GSD109D(FRD) |  |
| . | 9:51227405 |  | EEF1A1 | 0.054 | 0 | 3 | GSD67D(SRD),GSD73D(FRD),GSD109D(FRD) |  |
| . | 9:51227411 | downstream | EEF1A1 | 0.464 | 0 | 26 | GSD35D(FRD),GSD36D(SRD),GSD50D(FRD),GSD67D(SRD),  GSD68D(NRS,PTS),GSD73D(FRD),GSD77D(NRS, TS),GSD79D(FRD),  GSD80D(NRS,PTS),GSD85D(SRD),GSD90(ARD),GSD91D(SRD),  GSD92D(FRD),GSD93(SRD),GSD94(ARD),GSD95D(FRD),GSD96D(SRD),  GSD97D(SRD),GSD99D(FRD),GSD101D(SRD),GSD102D(SRD),  GSD103D(FRD),GSD104D(ARD),GSD105D(ARD),GSD108D(SRD),  GSD109D(FRD) |  |
| . | 9:51227774 | missense | EEF1A1 | 0.018 | 0 | 1 | GSD67D(SRD) |  |
| Variation | **Location** | **Consequence** | **Symbol** | **Maf** | **No call** | **Variant** | **Het** | **Homo** |
| . | 9:51227775 | stop_gained | EEF1A1 | 0.018 | 0 | 1 | GSD67D(SRD) |  |
| . | 9:51228186 | stop_gained | EEF1A1 | 0.464 | 0 | 26 | GSD35D(FRD),GSD36D(SRD),GSD50D(FRD),GSD67D(SRD),  GSD68D(NRS,PTS),GSD73D(FRD),GSD77D(NRS,PTS),GSD79D(FRD),  GSD80D(NRS,PTS),GSD85D(SRD),GSD89D(NRS,PTS),GSD90D(ARD),  GSD91D(SRD),GSD92D(FRD),GSD93D(SRD),GSD94D(ARD),  GSD96D(SRD),GSD97D(SRD),GSD99D(FRD),GSD101D(SRD),  GSD102D(SRD),GSD104D(ARD),GSD105D(ARD),GSD108D(SRD),  GSD109D(FRD),GSD110D(SRD) |  |
| . | 9:51228274 | missense | EEF1A1 | 0.018 | 0 | 1 | GSD103D(FRD) |  |
| rs24570325 | 9:51228828 | upstream | EEF1A1 | 0.263 | 9 | 7 | GSD35D(FRD),GSD79D(FRD),GSD89D(NRS,PTS),GSD93D(SRD) | GSD97D(SRD), GSD104D(ARD),  GSD109D(FRD) |
| rs24570326 | 9:51228829 | upstream | EEF1A1 | 0.263 | 9 | 7 | GSD35D(FRD),GSD79D(FRD),GSD89D(NRS,PTS),GSD93D(SRD) | GSD97D(SRD), GSD104D(ARD),  GSD109D(FRD) |
| . | 9:51235491 | downstream | Novel gene | 0.024 | 7 | 1 | GSD85D(SRD) |  |
| . | 9:51235495 | downstream | Novel gene | 0.024 | 7 | 1 | GSD85D(SRD) |  |
| . | 9:51255528 | upstream | Novel gene | 0.038 | 2 | 2 | GSD97D(SRD),GSD103D(FRD) |  |
| . | 9:51255530 | upstream | Novel gene | 0.038 | 2 | 2 | GSD97D(SRD),GSD103D(FRD) |  |
| . | 9:51255537 | upstream | Novel gene | 0.043 | 5 | 2 | GSD97D(SRD),GSD103D(FRD) |  |
| . | 9:51255622 | upstream | Novel gene | 0.232 | 0 | 11 | GSD35D(FRD),GSD50D(FRD),GSD67D(SRD),GSD89D(NRS,PTS),  GSD90D(ARD),GSD91D(SRD),GSD94D(ARD),GSD99D(FRD),  GSD109D(FRD) | GSD97D(SRD),  GSD103D(FRD) |
| Variation | **Location** | **Consequence** | **Symbol** | **Maf** | **No call** | **Variant** | **Het** | **Homo** |
| . | 9:51307901 | upstream | RALGDS | 0.018 | 0 | 1 | GSD99D(FRD) |  |
| . | 9:51330743 | downstream | RALGDS | 0.018 | 0 | 1 | GSD94D(ARD) |  |
| . | 9:51330772 | downstream | CEL | 0.018 | 0 | 1 | GSD94D(ARD) |  |
| . | 9:51344561 | downstream | GTF3C5 | 0.018 | 0 | 1 | GSD94D(ARD) |  |
| rs851223292 | 9:51345179 | missense | CEL | 0.018 | 0 | 1 | GSD94D(ARD) |  |
| . | 9:51403997 | upstream | MDH2 | 0.018 | 0 | 1 | GSD94D(ARD) |  |
| . | 9:51404393 | missense | MDH2 | 0.089 | 0 | 5 | GSD68D(NRS,PTS),GSD91D(SRD),GSD93D(SRD),GSD95D(FRD),  GSD103D(FRD) |  |
| . | 9:51404400 | missense | MDH2 | 0.107 | 0 | 6 | GSD67D(SRD),GSD85D(SRD),GSD95D(FRD),GSD102D(SRD),  GSD105D(ARD),GSD110D(SRD) |  |
| . | 9:51404429 | missense | MDH2 | 0.071 | 0 | 4 | GSD50D(FRD),GSD67D(SRD),GSD73D(FRD),GSD91D(SRD) |  |
| . | 9:51404724 | missense | MDH2 | 0.429 | 7 | 18 | GSD36D(SRD),GSD50D(FRD),GSD67D(SRD),GSD68D(NRS,PTS),  GSD77D(NRS,PTS),GSD79D(FRD),GSD80D(NRS,PTS),  GSD85D(NRS,PTS),GSD90D(ARD),GSD91D(SRD),GSD96D(SRD),  GSD97D(SRD),GSD99D(FRD),GSD101D(SRD),GSD102D(SRD),  GSD104D(ARD), GSD105D(ARD),GSD110D(SRD) |  |
| . | 9:51404910 | downstream | MDH2 | 0.018 | 0 | 1 | GSD50D(FRD) |  |
| . | 9:51404917 | downstream | MDH2 | 0.018 | 0 | 1 | GSD103D(FRD) |  |
| . | 9:51405007 | downstream | MDH2 | 0.054 | 0 | 1 | GSD103D(FRD) |  |

| Variation | Location | Consequence | Symbol | Maf | No call | Variant | Het | Homo |
| --- | --- | --- | --- | --- | --- | --- | --- | --- |
| . | 9:51405189 | downstream | MDH2 | 0.479 | 4 | 23 | GSD35D(FRD),GSD50D(FRD),GSD67D(SRD),GSD73D(FRD),  GSD77D(NRS, PTS),GSD79D(FRD),GSD80D(NRS,PTS),GSD85D  (SRD), GSD89D(NRS,PTS),GSD91D(SRD),GSD92D(FRD),GSD94D  (ARD),GSD95D(FRD),GSD96D(SRD),GSD97D(SRD),GSD99D(FRD),  GSD101D(SRD),GSD102D(SRD),GSD103D(FRD), GSD104D(ARD),  GSD105D(ARD),GSD108D(SRD),GS110D(SRD) |  |
| . | 9:51900804 | Upstream | SETX | 0.018 | 0 | 1 |  | GSD102D(SRD) |
| rs852815996 | 9:51994360 | downstream | NTNG2 | 0.55 | 8 | 12 | GSD102D(SRD),GSD103D(FRD) | GSD67D(SRD),  GSD73D(FRD),  GSD77D(NRS, PTS),  GSD79D(FRD),  GSD85D(SRD),  GSD94D(ARD),  GSD95D(FRD),  GSD104D(ARD),  GSD105D(ARD),  GSD108D(SRD) |

Variants on Chr11 and cases carrying each variant.

| Variation | Location | Consequence | Symbol | Maf | No call | Variant | Het | Homo |
| --- | --- | --- | --- | --- | --- | --- | --- | --- |
| . | 11:19522896 | downstream | pseudogene | 0.167 | 16 | 3 | GSD80D(NRS,PTS),GSD97D(SRD) | GSD108D(SRD) |
| . | 11:19522949 | downstream | pseudogene | 0.037 | 1 | 2 | GSD109D(FRD), GSD67D(SRD) |  |
| . | 11:20074787 | missense | ATP5MF | 0.5 | 26 | 1 |  | GSD36D(SRD) |
| . | 11:20143209 | downstream | MEIKIN | 0.04 | 3 | 1 |  | GSD105D(ARD) |
| rs851236833 | 11:20329483 | Upstream | IL3 | 0.280 | 3 | 12 | GSD35D(FRD),GSD50D(FRD),GSD79D(FRD),  GSD99D(FRD),GSD104D(ARD),GSD105D(ARD), GSD109D(FRD),GSD68D(NRS,PTS),GSD97D(SRD),  GSD101D (SRD) | GSD94D(ARD),  GSD67D(SRD) |
| rs853055449 | 11:20329514 | Upstream | IL3 | 0.280 | 3 | 12 | GSD35D(FRD),GSD50D(FRD),GSD79D(FRD),  GSD99D(FRD),GSD105D(ARD),GSD109D(FRD),  GSD93D(SRD),GSD96D(SRD),  GSD97D(SRD), SD101D(SRD) | GSD104D(ARD),  GSD67D(SRD) |
| rs852084335 | 11:20549348 | Upstream | P4HA2 | 0.250 | 2 | 12 | GSD35D(FRD),GSD50D(FRD),GSD79D(FRD,GSD90D (ARD),GSD99D(FRD),GSD103D(FRD),GSD104D(ARD),  GSD109D(FRD), GSD68D(NRS,PTS), GSD93D  (SRD), GSD101D(SRD) | GSD94D(ARD) |
| rs850644440 | 11:20587679 | downstream | PDLIM4 | 0.316 | 9 | 9 | GSD35D(FRD),GSD50D(FRD),GSD77D(NRS,PTS),  GSD103D(FRD), GSD105D(ARD), GSD109D(FRD) | GSD67D(SRD),GSD79D  (FRD), GSD104D(ARD) |
| . | 11:20597980 | Upstream | SLC22A4 | 0.071 | 0 | 2 |  | GSD105D(ARD),  GSD95D(FRD) |
| rs22191909 | 11:20908625 | downstream | pseudogene | 0.018 | 0 | 2 | GSD95D(FRD) | GSD108D(SRD) |
| . | 11:20909122 | downstream | pseudogene | 0.105 | 9 | 2 |  | GSD68D(NRS,PTS), GSD101D(SRD) |

| Variation | Location | Consequence | Symbol | Maf | No call | Variant | Het | Homo |
| --- | --- | --- | --- | --- | --- | --- | --- | --- |
| . | 11:20909130 | downstream | pseudogene | 0.105 | 9 | 2 |  | GSD68D (NRS,PTS), GSD101D(SRD) |
| . | 11:20909131 | downstream | pseudogene | 0.263 | 9 | 6 | GSD36D(SRD), GSD97D(SRD) | GSD68D(NRS,PTS),GSD92D(FRD),GSD99D  (FRD), GSD101D(SRD) |
| rs22191752 | 11:20962569 | downstream | IL13 | 0.423 | 2 | 14 | GSD50D(FRD),GSD73D(FRD), GSD90D(ARD),GSD92D(FRD), GSD99D(FRD),GSD109D FRD) | GSD36D(SRD),GSD79D(FRD),GSD93D(SRD), GSD94D(ARD),GSD97D(SRD),GSD103D(FRD), GSD104D(ARD), GSD105D(ARD) |
| . | 11:20982595 | downstream | IL4 | 0.13 | 1 | 5 | GSD79D(FRD),GSD105D(ARD), GSD97D(SRD), | GSD92D (FRD),GSD36D(SRD) |
| . | 11:21044750 | Upstream | KIF3A | 0.018 | 0 | 1 | GSD97D (SRD) |  |
| . | 11:21051992 | downstream | SEPTIN8 | 0.13 | 1 | 5 | GSD93D(SRD),GSD95D(FRD), GSD97D(SRD) | GSD36D(SRD), GSD105D(ARD) |
